# Temporally-Divergent Regulatory Mechanisms Govern Neuronal Development and Diversification in the Neocortex

**DOI:** 10.1101/2020.08.23.263434

**Authors:** Wen Yuan, Sai Ma, Juliana R. Brown, Kwanho Kim, Vanessa Murek, Lucia Trastulla, Alexander Meissner, Simona Lodato, Ashwin Shetty, Joshua Z. Levin, Jason D. Buenrostro, Michael J. Ziller, Paola Arlotta

## Abstract

Mammalian neocortical neurons span one of the most diverse cell type spectra of any tissue. The regulatory strategies that neurons use during progressive development and maturation remain unclear. We present an integrated single-cell epigenomic and transcriptional analysis of individual classes of neurons from both mouse and marmoset neocortex, sampled during both early postmitotic stages of identity acquisition and later stages of neuronal plasticity and circuit integration. We find that in both species, the regulatory strategies controlling these early and late stages diverge: early postmitotic neurons use molecular regulatory programs with broader tissue distribution and greater evolutionary conservation, while programs active during later neuronal maturation implement more brain- and neuron-specific mechanisms showing greater evolutionary divergence. The data uncovers a temporally-regulated shift in regulatory choices, likely reflecting unique evolutionary constraints on distinct events of neuronal development in the neocortex.

## Introduction

The mammalian cerebral cortex contains a great diversity of neurons, which differ in their connectivity, neurotransmitter usage, morphology, gene expression, and electrophysiological properties (Lodato and Arlotta, 2015). Although recent work has uncovered the molecular states that define different classes of terminally differentiated neurons in the adult brain (Darmanis et al., 2015; Gray et al., 2017; Lake et al., 2016; Mo et al., 2015; Tasic et al., 2016; Yao et al., 2020; Zeisel et al., 2015), profiling of cortical neurons over developmental time courses has been limited and largely confined to transcriptional profiling of a few embryonic and neonatal stages (Frazer et al., 2017; Molyneaux et al., 2015), or of single adult timepoints (Cusanovich et al., 2018; Frazer et al., 2017; Hodge et al., 2019; Lake et al., 2018; Luo et al., 2017; Nott et al., 2019). There is not yet a holistic picture of the molecular dynamics and regulatory landscapes of individual cortical neuron classes, or any other class of mammalian central neurons, over extended trajectories of cell identity acquisition and maturation during postnatal life. This has precluded in-depth understanding of the molecular strategies used by neurons to acquire their identity, mature, and wire. Similarly, how such molecular strategies might vary across species has not been addressed.

Here, we defined transcriptional and epigenomic changes in different populations of postmitotic cortical neurons over a timeline spanning perinatal stages to adulthood in two species, mouse and marmoset, to elucidate the regulatory strategies employed over progressive stages of postmitotic development and maturation.

## Results

### Early and late stages of cortical pyramidal neuron postmitotic development use divergent regulatory programs

Even after becoming postmitotic, cortical neurons undergo extensive development, from establishment of subtype identity to postnatal refinement of terminally differentiated features. It is largely not understood whether regulatory strategies at play during postmitotic development remain constant over postnatal life.

To understand these regulatory strategies, we first applied inducible Cre mouse lines to examine two major classes of neocortical pyramidal neurons. We profiled *Cux2*-lineage layer 2/3 (L2/3) callosal projection neurons (CPNs), which are involved in associative functions and are the most recently-evolved population of cortical neurons (henceforth referred to as “Cux2 CPNs”), and *Tle4*-lineage layer 6 (L6) corticothalamic projection neurons (CThPNs), which are responsible for integration of sensory and motor information (henceforth referred to as “Tle4 CThPNs”). Each type was isolated across a timecourse spanning establishment of class-specific neuronal identity and early neuronal development (embryonic day 18.5 [E18.5] and postnatal day 1 [P1], P3, and P7), periods of cortical plasticity (P21), and neuronal maturation and integration into cortical circuits (P21 and P48) (Fig. 1a and Fig. S1, S2 and supplementary text). Labelled neurons from dissected somatosensory cortex were isolated by fluorescence-activated cell sorting (FACS), and profiled in bulk for gene expression by RNA sequencing (RNA-seq), for DNA methylation (DNAme) by whole-genome bisulfite sequencing (WGBS), and for open chromatin by assay for transposase-accessible chromatin using sequencing (ATAC-seq).

**Fig. 1.**
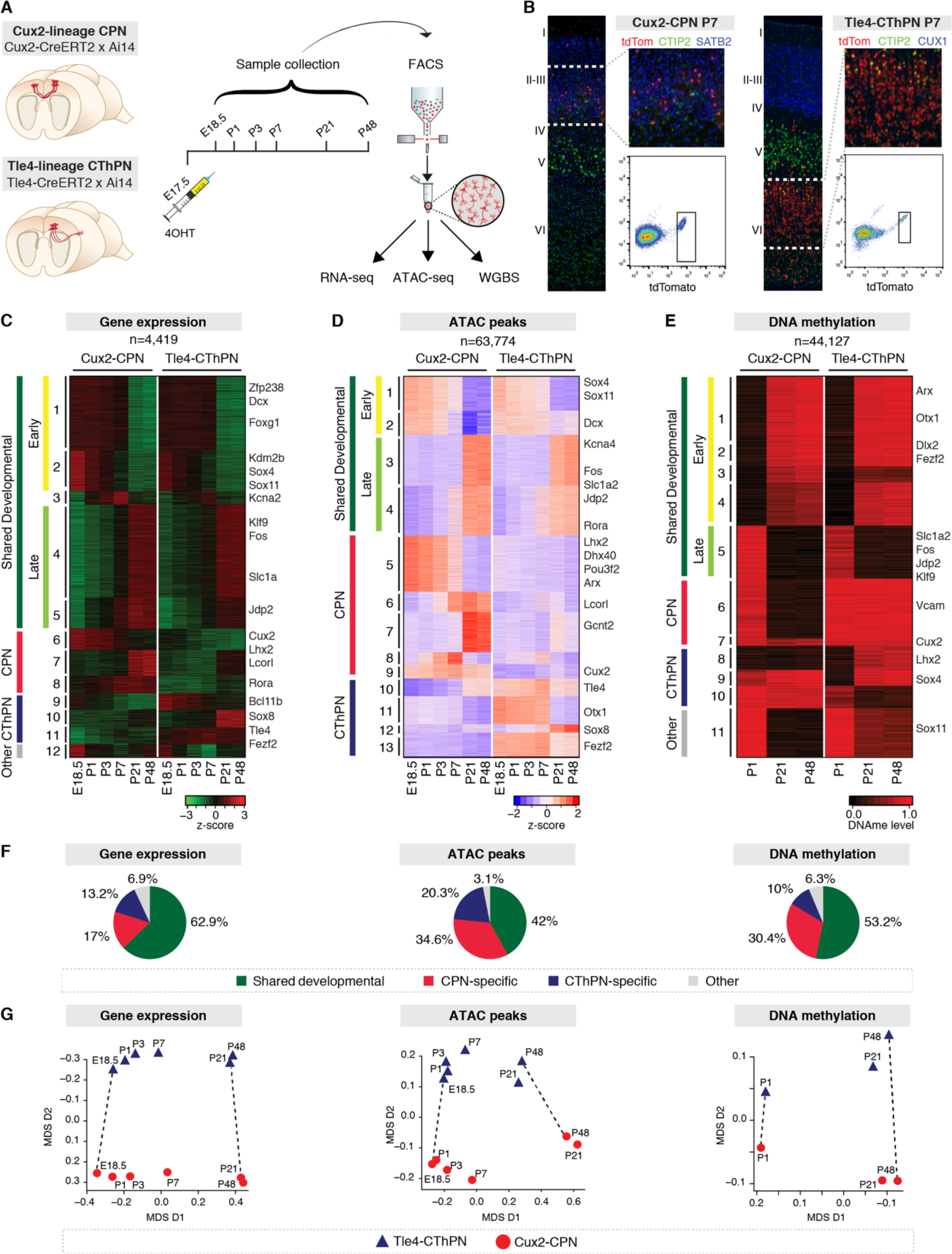
Profiling of genetically-defined cortical pyramidal neuron classes. (**A**) Schematic of experimental design. (**B**) Representative coronal sections showing correct laminar location of tdTomato^+^ cells in the somatosensory cortex and FACS plots of Tle4-CThPNs and Cux2-CPNs at P7. Also see Fig. S1. (**C-E**) Developmental dynamics of differentially expressed genes (**C),** differentially enriched ATAC-seq peaks (**D)**, and differentially methylated regions (DMRs) (**E)**. (**F)** Fraction of dynamic features classified into each overall category. (**G**) 2D MDS plots of dynamic feature sets over time for each dataset.

We identified features (transcribed genes, differentially accessible chromatin peaks, and differentially-methylated regions [DMRs]) that were dynamic over age or between cell types (examples in Fig. S3), and applied k-means clustering to group features with similar patterns (Fig. 1C-E and Fig. S4). We find that for all datasets, 40-60% of dynamic features were assigned to clusters that were associated with developmental stage and independent of neuronal subtype (Fig. 1C-F). These shared, developmentally-regulated clusters fell into two major categories: those predominantly active (transcriptionally upregulated, accessible, or hypomethylated) at embryonic and early postnatal ages (E18.5-P7; “early developmental”, yellow bars in Fig. 1C-E), and those predominantly active at weaning and older ages (P21-P48; “late developmental”, light green bars in Fig. 1C-E).

A smaller proportion of dynamic features showed neuron-class-specific patterns; as expected, these clusters included known molecular markers of CPNs and CThPNs (Arlotta et al., 2005; Galazo et al., 2016) (Fig. 1C-E). Notably, while class-specific clusters accounted for only 23% of dynamic transcriptional features, they comprised 34% (DNAme) to 45% (ATAC) of dynamic epigenetic features (Fig. 1F), indicating higher neuronal subtype-specificity of epigenetic changes, in agreement with findings that epigenetic signatures may be particularly powerful in discriminating neuronal subclasses (Luo et al., 2017; McKinley et al., 2020).

Visualization using multi-dimensional scaling (MDS), which preserves relative distance between datapoints, showed that, despite widespread changes over development, these two classes of neurons did not change in overall similarity with time in gene expression or open chromatin profiles (Fig. 1G and Fig. S4B-C). In contrast, the DNA methylation landscape becomes more divergent between neuronal subtypes over postnatal life (p<0.05, two-sided t-test; Fig. 1G and Fig. S4C); this increase was not driven by global changes in methylation or expression of DNA methylases (Fig. S5A-B), but rather by changes in distribution patterns across the genome. DNA methylation increased over time at genes and gene regulatory elements (GREs, inferred from open chromatin sites) characteristic of other cell types and of earlier developmental stages (Fig. S5C), consistent with its known role in stabilizing silencing of transcriptional programs associated with other fates or developmental stages (Guibert and Weber, 2013).

We investigated the strategies for genome regulation that are common to pyramidal neurons across developmental time, by examining clusters whose temporal dynamics were shared between both cell types (“shared early” and “shared late” clusters, Fig 1C-E). Detailed characterization of these different feature sets uncovered a striking temporal transition in regulatory principles, across multiple modalities, between shared programs of early (E18.5-P7) vs. late (P21-P48) postmitotic neuronal development.

First, we investigated the expression of both all genes in each cluster and of transcription factors (TFs) across 77 mouse brain regions and developmental timepoints from the Allen Brain Atlas (Thompson et al., 2014) and 294 mouse cell types and tissues from the FANTOM5 project (FANTOM Consortium and the RIKEN PMI and CLST (DGT) et al., 2014) (Fig. 2 B-C and S6A). We found that for both all genes and for TFs only, the shared late-active genes showed significantly more temporal and/or cell-type specificity of expression in both tissue panels (p ≤ 2.576 − 10^−7^ for all comparisons, one-sided Mann-Whitney Test), suggesting that early gene and TF programs reflect more “generic” developmental processes.

**Fig. 2.**
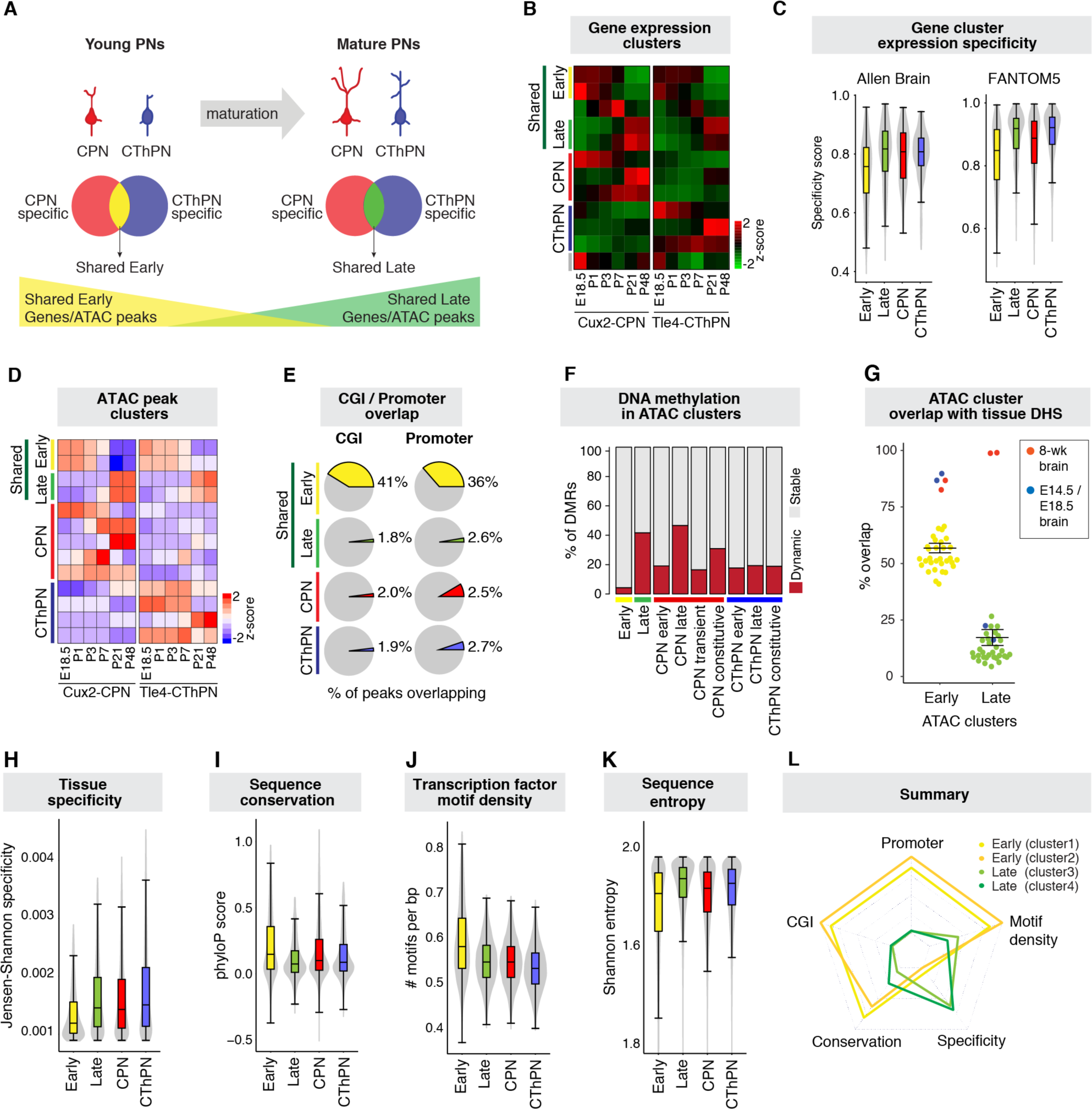
Divergent regulatory principles of early and late stages of neuronal development. (**A**) Schematic of analyses. (**B**) Summary of gene expression clusters from Fig. 1C. (**C**) Transcription factor expression specificity within the brain for TFs in different categories of cluster, from expression data from the Allen Brain Atlas (left). Higher values indicate more specific expression. Transcription factor expression specificity across 397 mouse cell types from the FANTOM5 project (right). (**D**) Summary of ATAC peak clusters from Fig. 1D. (**E**) Fraction of open chromatin regions (ATAC-seq peaks) in different categories of clusters that overlap CpG islands (CGI) or annotated promoter regions. (**F**) Fraction of DMRs in ATAC clusters of different categories which are static or dynamic over the timecourse. (**G**) Fraction of ATAC-seq peaks from different classes of ATAC clusters that overlap DNAse hypersensitivity sites across 35 cell types from the mouse ENCODE project. Adult (8-week) whole-brain and telencephalon samples highlighted in red; embryonic (E14.5 and E18.5) brain samples highlighted in blue. Also see Fig. S6F. (**H**) Open chromatin specificity across ATAC peak clusters, as the concordance of ATAC peaks with a scATAC-seq panel of 85 mouse primary tissues and cell types (Cusanovich et al., 2018). (**I**) Distribution of phyloP sequence conservation across all placental mammals for early- and late-developmental ATAC clusters. (**J**) Density of known transcription factor binding motifs within ATAC clusters of different types. (**K**) Average sequence entropy within ATAC clusters of different types. (**L**) Summary of characteristics from C and G-J for each of the shared developmental ATAC peak clusters individually. Also see Fig. S6B.

We then characterized putative GREs identified from our ATAC-seq profiles, and found that GREs in shared early-developmental ATAC clusters are more than ten-fold enriched for annotated promoter regions (within 1 KB up- or downstream of the TSSs) and CpG islands (Fig. 2D-E and Fig. S6B), which are strongly associated with TSSs and play an important role in gene regulation (Deaton and Bird, 2011). Conversely, members of the shared-late developmental ATAC clusters are largely TSS-distal and show lower frequency of overlap with CpG islands and annotated promoters (Fig. 2D-E and Fig. S6B), suggesting that they disproportionately function as distal regulatory elements such as enhancers or insulators. This division indicates that different regulatory strategies (promoter-based vs. enhancer/insulator-based) underlie early events of subtype identity acquisition vs. later phases of neuronal plasticity and integration. Interestingly, cell-type-specific ATAC clusters showed low overlap with CpG islands and annotated promoters regardless of developmental dynamics (Fig. 2D-E and Fig. S6B).

To investigate the relationship between the shared early- vs late-active gene expression and ATAC peak clusters, we calculated the overlap between genes in the RNA-seq clusters and putative genes regulated by GREs in the ATAC clusters (defined as peaks within 100 kb of TSSs). We found that the early gene expression clusters were most strongly correlated with the early ATAC clusters, and similarly for the late gene expression and late ATAC clusters (Fig. S6C), indicating that the shared developmental transcriptional and epigenetic programs affect similar sets of genes.

We next investigated the relationship between changes in chromatin state and DNAme state across cell types and developmental time. This analysis showed a significant overlap between clusters with similar dynamics, where regions losing open chromatin gained DNAme and vice versa, for shared early, shared late, and cell type-specific clusters (Fig S6E). However, the fraction of open chromatin regions exhibiting temporally-dynamic DNAme patterns was significantly higher in the shared-late ATAC cluster compared to the shared-early cluster (41.72% vs. 4.29%; Fig. 2F and Fig. S6E). This observation is in line with the elevated frequency of promoter and CpG island (CGI) regions in the shared early clusters, and the much higher frequency of distal putative regulatory elements in the shared late clusters (Fig. 2D-E, Fig. S6B). Promoter regions are well known to be much less prone to DNAme changes over development, whereas distal differentially accessible sites frequently coincide with enhancer regions and transcription factor binding sites that exhibit highly dynamic DNAme patterns (Ziller et al., 2013). This observation suggests that DNA methylation plays a greater role in regulating late developmental programs by mediating stable silencing of distal putative regulatory elements.

Recent studies have suggested that cell-type-specific accessible chromatin sites are preferentially localized in putative enhancer regions compared to promoter regions (Corces et al., 2016; Gray et al., 2017; Natarajan et al., 2012; Song et al., 2011). We therefore evaluated the tissue specificity of early-active vs. late-active GREs across a panel of DNase hypersensitivity sites in 35 adult and embryonic mouse primary tissues and cell types from the ENCODE database (Yue et al., 2014). Strikingly, while GREs in shared early-developmental epigenetic clusters frequently resided in an open chromatin state in many tissues, GREs in shared late-developmental epigenetic clusters were on average found in an open state in significantly fewer tissues (ATAC clusters early vs late: p < 2.2 × 10^−16^ Mann-Whitney test; Fig. 2G and Fig. S6F). Most notably, late-active ATAC clusters are highly enriched for chromatin regions that are accessible in adult (8 week) mouse brain tissues, but not for those open in embryonic brain, adult cerebellum, adult and perinatal retina, or any non-CNS tissue. In contrast, GREs in early-active ATAC clusters show high overlap with adult and embryonic brain but also show broad associations with open chromatin regions in many other cell types and tissues (Fig. 2G, Fig. S6F). The ratio of accessible sites suggests that the shared late-active regions are highly enriched for GREs that are specific to the adult brain, consistent with the large number of genes characteristic of neuron-specific processes in these clusters (Table S1-2).

Collectively, these data suggest that genes and gene regulatory features (TFs and GREs) that are active in early-postmitotic stages of pyramidal neuron development are comparatively more widely used across many tissues and cell types, and therefore may represent more “generic” basic developmental programs, while those active at later stages are more cell-type specific, and in particular characteristic of the adult brain, and thus may control more specialized processes in nervous system function.

Given these findings, we examined further potential differences between the shared early and late ATAC regions. We first compared the average tissue specificity of accessible sites across a scATAC-seq panel of 85 mouse primary tissues and cell types (Cusanovich et al., 2018) (see Methods). Early-active sites showed lower average specificity (Fig. 2H), again indicating that these sites are more widely used across tissues.

Previous analyses of DNA methylation have reported that sequence conservation varies for DMRs characteristic of different tissue types, with DMRs characteristic of ectodermal tissues showing highest conservation (Hon et al., 2013), and that DMRs specific to excitatory cortical neurons are less conserved than those specific to interneurons (Luo et al., 2017). Given these observations and the more ubiquitous activity across tissues of genes and GREs in the shared early-developmental clusters, we hypothesized that these regions might be under different degrees of evolutionary constraint. We thus investigated the evolutionary conservation across placental mammals of the ATAC peak regions in the shared developmentally-regulated ATAC clusters, and detected significantly higher sequence conservation of elements open at early stages (p < 2.2 − 10^−16^, Mann-Whitney Test) (Fig. 2I and Fig. S6B). Consistent with this finding, shared early-active GREs also showed a higher density of TF binding motifs (Fig. 2J), and lower sequence entropy (Fig. 2K), a metric of sequence constraint, both potentially associated with increased CpG density. Interestingly, across all of these metrics, the neuron subtype-specific gene and GRE clusters largely resembled the late shared-developmental clusters (Fig. 2B-E and H-L, and Fig. S6B).

Together, these findings (summarized in Fig. 2L) suggest that early-developmental GREs may use more widely-shared regulatory mechanisms that are consequently under greater evolutionary constraint. In contrast, late-developmental GREs, which have more restricted tissue usage, may be more amenable to variation, and may employ more species-specific or evolutionarily recent mechanisms.

### Temporal divergence in global regulatory strategies is a general principle of cortical neuron maturation across species

Next, we sought to investigate whether these regulatory principles are common to all neuronal classes in the cortex, and whether cortical neurons in non-human primates follow the same principles as in mouse, using marmoset as a model system. We performed single-cell (mouse) or single-nucleus (marmoset) RNA sequencing (scRNA-seq, snRNA-seq) and single-nucleus (mouse) or single-cell (marmoset) ATAC sequencing (snATAC-seq, scATAC-seq) on unfractionated cortical tissue from these two species across early and late stages of postnatal development. We profiled combined somatosensory and motor cortex from mice at three selected ages: P1, representing a period when early-active gene and GRE clusters predominate, P7, as an intermediate stage, and P21, representing a period when late-active gene and GRE clusters predominate. Marmoset somatosensory cortex was profiled at neonatal (P1-P2) and adult ages (2 years [Y2] for scRNA-seq, Y7-8 for scATAC-seq, based on limited tissue availability for this species).

We first examined programs of gene expression across the two species. After quality control and filtering, the final dataset contained a total of 60,989 mouse and 36,592 marmoset cells across the different timepoints (Fig. 3A-B; Fig. S7-10). To identify individual cell populations within the dataset, we performed cell clustering on the sc/snRNAseq data and assigned cell identity based on expression of known canonical marker genes (Fig. 3A-D; Fig. S8; Fig. S10). We identified major neuronal and glial cell populations, including different populations of pyramidal neurons (CPN, subcerebral projection neurons [SCPN], and CThPN) and interneurons, astrocytes, and oligodendrocyte-lineage populations. Fig. 3C-D shows expression of example classical marker genes for selected populations, including Neurod2 and TBR1 (glutamatergic neurons), Gad2 (GABAergic interneurons), Pdgfra (oligodendrocyte lineage), and Aqp4 (astrocytes). For both mice and marmoset, clusters showed separation by both age and cell type, with projection neuron populations being predominantly separated by age (Fig. 3A-B; Fig. S7C; Fig. S9C), consistent with our findings that the majority of changes in the mouse Cux2-CPN/Tle4-CThPN bulk sequencing datasets are developmentally-related rather than neuron subtype-specific (Fig 1).

**Fig. 3.**
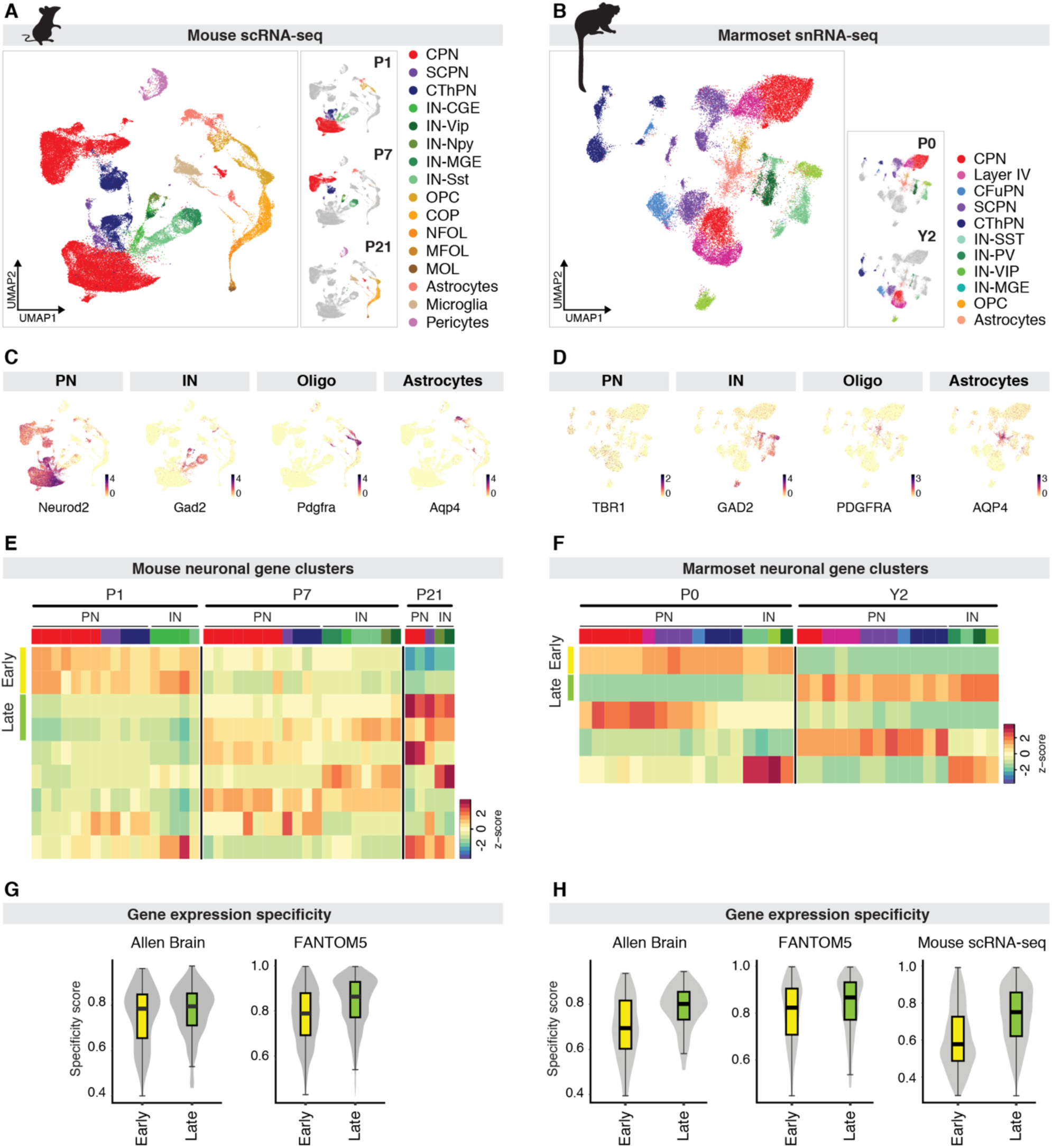
Single-cell RNA sequencing demonstrates a developmental shift in specificity of shared gene expression programs across multiple neuronal subclasses in both mouse and marmoset. (**A**) UMAP representation of gene expression profiles from 60,989 single cells from mouse cortex at P1, P7, and P21, color-coded by major cell type. Left: UMAP plots showing cell distribution by age. (**B**) UMAP representation of 36,592 single nuclei from marmoset cortex at P0 and 2 years (Y2), color-coded by major cell type. Left: UMAP plots showing cell distribution by age. (**C**) Representative marker genes for major cell types in the mouse data. Also see Fig. S8 A, C. (**D**) Representative marker genes for major cell types in the marmoset data. Also see Fig. S10 A, C. (**E**) Developmental dynamics of clusters of differentially expressed genes across the mouse excitatory and inhibitory neuronal populations (cell type indicated by color-coded bar at top, corresponding to colors in A). (**F**) Developmental dynamics of clusters of differentially expressed genes across the marmoset excitatory and inhibitory neuronal populations (cell type indicated by color-coded bar at top, corresponding to colors in B). (**G**) Mouse gene expression specificity in the shared-early and shared-late gene clusters, within the brain (from expression data from the Allen Brain Atlas; left), and across 397 mouse cell types (from the FANTOM5 project; right). Higher values indicate more specific expression. (**H**) Marmoset gene expression specificity in the shared-early and shared-late gene clusters, within the brain (from expression data from the Allen Brain Atlas; left), across 397 mouse cell types (from the FANTOM5 project; center), and across cell populations from our mouse single-cell dataset (right). Higher values indicate more specific expression.

We then identified the shared-developmental and cell type-specific gene clusters. For each species, we selected all pyramidal neuron and interneuron populations, identified genes that showed differential expression across the sc/snRNA-seq datasets, and clustered them by their expression pattern across age and cell type as we did for the bulk pyramidal neuron sequencing data (Fig. 3E-F). For mice, we identified two early and two late shared developmentally-regulated gene clusters which were shared across all pyramidal and interneuron populations (indicated by yellow bars, Fig. 3E-F), and for marmoset we identified one early and one late pan-neuronal gene cluster (indicated by light green bars). We repeated the tissue- and cell type-specificity analyses for genes and TFs in these clusters, and found that, for both species, the late developmental clusters showed greater specificity across both the Allen (brain) and FANTOM5 (tissue) datasets (Fig. 3G-H and Fig. S11B, D). In addition, the marmoset shared late gene cluster showed greater cell type-specificity across our mouse single-cell RNA-seq dataset (Fig. 3H). Notably, although the pyramidal and interneuron cell types could be further clustered, both separating different sub-classes and further dividing some subclasses, we did not observe striking differences within the subclusters of individual types, indicating that heterogeneity within neuron populations does not explain the gene clustering results. The data indicate that the shift in global regulatory principles observed in CPN and CThPN applies to all cortical neuron subtypes, and to non-human primate cortex.

Next, to examine regulatory characteristics of GREs across cell types and species, we performed single-cell (sc) or single-nucleus (sn) ATAC-seq at neonatal and juvenile/adult ages for both species (Fig. 4A-B and see Methods). After quality control and filtering, the final dataset included 19,145 mouse and 15,919 marmoset cells/nuclei (Fig. 4A-B; Fig. S12-S14). Cell-type identities were assigned to cell clusters based on inferred expression of the same panel of known marker genes used in the sc/snRNA-seq analyses (Fig. S14A-D). Example gene tracks of selected canonical marker genes across each cell cluster are shown in Fig. S14B,D. Similar to the scRNA-seq data, neuron populations were predominantly separated by age in both species (Fig. S12B and Fig. S13B). We then collapsed differentially-accessible ATAC peak regions by cell cluster and clustered these pseudo-bulk profiles. We identified early-active and late-active clusters shared across pyramidal neuron and interneuron classes (Fig. 4C, E). For mice, we identified one pan-neuronal early peak cluster and one pan-neuronal late peak cluster, as well as two late peak clusters shared across pyramidal neuron populations but not interneurons (Fig. 4C).

**Fig. 4.**
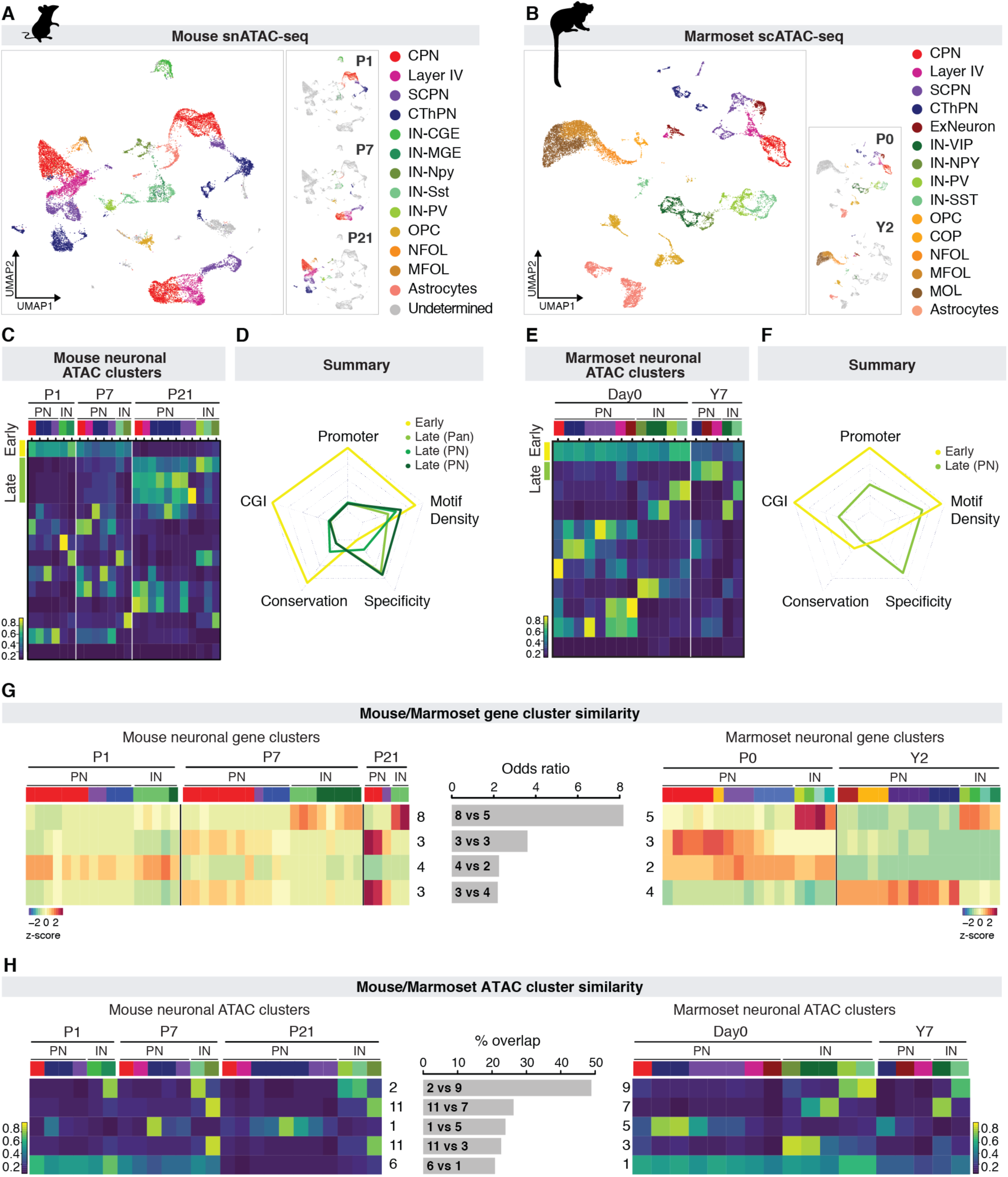
Developmental shift in gene regulatory principles is common to multiple neuronal subclasses and is conserved across mouse and marmoset. (**A**) UMAP representation of ATAC chromatin accessibility profiles from 19,145 single nuclei from mouse cortex at P1, P7, and P21, color-coded by major cell type. Left: UMAP plots with cell distribution by age. (**B**) UMAP representation of ATAC chromatin accessibility profiles from 15,919 single cells from marmoset cortex at P0 and 2 years (Y2), color-coded by major cell type. Left: UMAP plots with cell distribution by age. (**C**) Developmental dynamics of clusters of differentially accessible ATAC peaks across the mouse excitatory and inhibitory neuronal populations (cell type indicated by color-coded bar at top, corresponding to colors in A). (**D**) Summary of CpG island overlap, promoter overlap, TF motif density, tissue specificity, and sequence conservation for each of the mouse shared developmentally-regulated clusters, as in Fig. 2K. Also see Fig. S15A-C. (**E**) Developmental dynamics of clusters of differentially accessible ATAC peaks across the marmoset excitatory and inhibitory neuronal populations (cell type indicated by color-coded bar at top, corresponding to colors in D). (**F**) Summary of CpG island overlap, promoter overlap, TF motif density, tissue specificity, and sequence conservation for each of the marmoset shared developmentally-regulated clusters. Also see Fig. S15D-F. (**G**) Overlap between genes in the mouse and marmoset single-cell gene expression clusters, showing the 4 most similar pairs. Also see Fig. S16A. (**H**) Overlap between accessible regions in the mouse and marmoset single-cell ATAC chromatin accessibility clusters, showing the 5 most similar pairs. Also see Fig. S16C.

Importantly, the dynamics of the scATAC peak clusters are correlated with DNAme dynamics (taken from the genetically-labelled, FACS purified CPN and CThPN dataset): early pan-neuronal scATAC peaks remain mostly unmethylated while late pan-neuronal peaks lose methylation over developmental time (Fig. S16E). Interestingly, glia-specific open chromatin regions (such as cluster 2 and 4) show relatively low DNAme levels in P1 CPNs and CThPNs, and only gain DNAme in these neurons during maturation (Fig. S16F), consistent with the progressive silencing of alternative lineage programs by DNAme (Ehrlich and Lacey, 2013). Interneuron-specific open chromatin regions (cluster 7), however, show consistently high DNAme levels in CPNs and CThPNs across all ages, suggesting that these programs may need to be silenced at an earlier stage in pyramidal neuron development.

For marmoset, we likewise identified one pan-neuronal early peak cluster, and one late peak cluster shared across pyramidal neurons but not interneurons. However, we did not identify a pan-neuronal late peak cluster (Fig. 4E). This may suggest that, in marmoset, adult regulatory programs may diverge more between interneurons and projection neurons than they do in mice, which may reflect greater neuronal specialization in more evolutionarily advanced cortices

We repeated the ATAC peak characterization previously performed on the mouse Cux2-CPN/Tle4-CThPN bulk sequencing dataset on the snATAC-seq datasets. Consistent with the bulk data, we found that for both species, the shared early-active peak clusters showed greater enrichment for promoters and CGI, increased density of binding motifs, lower tissue specificity, and higher sequence conservation (Fig. 4D, F and Fig. S15C-D). Similar to the sc/snRNA-seq data, we did not observe significant differences within the neuronal subclusters in the shared developmental programs.

Together, these data show that the transition in shared developmental programs between broadly-used, more highly conserved regulatory elements at earlier stages of postmitotic development vs. more cell type- and tissue-specific regulatory elements at later stages holds true across neuronal classes, and is conserved in non-human primate cortex. The data suggests that these regulatory principles are broadly generalized properties of neuronal maturation.

To compare the degree of similarity between the shared-developmental gene and GRE programs between mouse and marmoset, we examined overlap between mouse and marmoset gene and ATAC peak clusters (Fig. 4G-H). We found that the shared-early gene clusters were among the most highly similar pairs of clusters; similarly, the shared-early ATAC peak clusters were among the most similar pairs between species (Fig. 4G-H). These findings indicate that general programs of early pan-neuronal development are more frequently shared between species compared to later pan-neuronal programs, which show greater species specificity. This observation is consistent with the broader sequence-level conservation found in both the bulk pyramidal neuron and single-nucleus ATAC-seq datasets. Interestingly, interneuron-specific clusters were the most highly similar between species for both the gene and ATAC-seq datasets, suggesting that interneuron-specific developmental programs are more highly conserved (Fig. 4G-H).

In sum, our analysis uncovered a striking temporal shift in regulatory principles, across multiple modalities, between shared programs of early (perinatal) and late (juvenile/adult) postmitotic neuronal development, that is shared between neuronal subtypes and species.

## Discussion

The neocortex contains a great diversity of neuronal classes, which are born during embryogenesis but undergo substantial postnatal maturation to acquire their adult features. Although these long trajectories of neuronal development are critical to build a functional cerebral cortex, the regulatory mechanisms that govern the execution of these sequential stages of postmitotic neuronal development and maturation are poorly understood.

Here we have defined a global outline of the regulatory principles underlying key steps of embryonic and postnatal development of cortical neurons in both rodents and non-human primates. In both species, we uncovered a striking developmental shift between two distinct strategies of epigenomic and transcriptional regulation. Early-active developmentally-shared programs (perinatal) employ more broadly-expressed transcription factors and more promoter-associated, tissue-nonspecific, and evolutionarily conserved regulatory sequences. Conversely, developmentally-shared programs in mature neurons (juvenile/adult) employ more tissue-restricted transcription factors and more distal, tissue- and cell-type specific (and indeed brain-specific), and evolutionarily divergent regulatory sequences. This shift applies to multiple neuronal classes and cannot be explained by variation within neuronal populations. Notably, these rules also apply across species, including non-human primates, suggesting that this temporal change in regulatory programs represents a broadly-applied, core strategy for cortical neuron development.

The data supports a conceptual framework in which fundamental events of cortical neuron development that occur during perinatal stages, such as establishment of neuronal identity and the acquisition of basic aspects of neuronal architecture, use molecular programs that are shared with other tissues, and are thus mediated by more “generic” regulatory programs, requiring a more-constrained degree of variation. It is tempting to speculate that this reflects the need for the nervous system to build its basic cell types in a reproducible and invariant manner. In contrast, as neurons transition to phases of neuronal, circuit, and synaptic plasticity and function, they employ more specialized transcriptional and epigenetic programs that may allow for more flexibility; the greater variation in cell and circuit behavior at later stages of cortical maturation may benefit from increased “customization”, reflected by more rapid species divergence in late developmental regulatory programs.

## Acknowledgments

We thank the laboratory of Guoping Feng for providing marmoset tissue samples. We thank Hsu-Hsin Chen for her advice and assistance in neuron isolation techniques. We thank Bruna Paulsen for assistance with preparing figures. We thank members of the Arlotta lab for insightful comments and suggestions. **Funding:** This work was supported by grants from NIH (R01MH101268, R01NS078164, and U19MH114821) to P.A, and by BMBF eMed program grant 01ZX1504 to M.J.Z. J.D.B. and the Buenrostro lab acknowledge support by the Allen Distinguished Investigator Program through the Paul G. Allen Frontiers Group, the Chan Zuckerberg Initiative, and support from the NIH New Innovator Award (DP2 mechanism).

## Supplementary Materials

### Methods

#### Mice

Experiments using mice were conducted under protocols approved by the Harvard University Institutional Animal Care and Use Committee and followed the guidelines set forth in the National Institute of Health Guide for the Care and Use of Laboratory Animals.

*Cux2*-lineage CPNs were labelled using a *Cux2*-CreERT2 knock-in line (NIH Neuroscience Blueprint Cre Driver Network, 2009). *Tle4*-lineage CThPNs were labelled with a *Tle4*-2A-CreERT2 mouse line (Matho et al., 2020). The CRE-inducible tdTomato reporter line Ai14 (Madisen et al., 2010) was used to detect recombined cells. *Cux2*-CreERT2;Ai14 or *Tle4*-2A-CreERT2;Ai14 double homozygous male mice were crossed with WT C57BL/6J females. Cre recombination was induced at E17.5, after the major wave of CPN neurogenesis (E15.5-E17.5) is complete (Fame et al., 2011). 4-Hydroxytamoxifen (4-OHT; Sigma) dissolved in corn oil was administered to pregnant mice at 1 mg 4-OHT/10g body weight.

For FACS isolation, CRE recombination was induced at E17.5 and the somatosensory cortex from transgenic animals was dissected and dissociated at E18.5, P1, P3, P7, P21 and P48. Tissue dissociation was performed as described (Arlotta et al., 2005), and td-Tomato-positive cortical pyramidal neurons were isolated by FACS. We performed RNA sequencing (RNA-seq) and assay for transposase-accessible chromatin using sequencing (ATAC-seq) at E18.5, P1, P3, P7, P21 and P48, and whole-genome bisulfite sequencing (WGBS) at P1, P21, and P48. Each library represents a pool of tissue from 10 animals (5 male and 5 female) from two litters. Two biological replicates were performed for each age and neuron type for each assay.

#### Marmoset

Marmoset tissue was obtained from the laboratory of Guoping Feng at the Massachusetts Institute of Technology. All marmoset experiments were approved by the Institutional Animal Care and Use Committee of Massachusetts Institute of Technology and followed the guidelines from the National Institute of Health’s Guide for the Care and Use of Laboratory Animals. For tissue collection, adult marmosets were deeply sedated with ketamine (20-40 mg/kg, IM) and/or alfaxalone (5-10 mg/kg, IM), followed by intravenous injection of sodium pentobarbital (10– 30 mg/kg). Because venous access was not possible in neonates, infant marmosets were sedated with intraperitoneal injection of sodium pentobarbital (10– 30 mg/kg). When pedal withdrawal reflex was eliminated and/or respiratory rate was diminished, animals were transcardially perfused with ice-cold PBS or Sucrose-HEPES buffer. Whole brains were rapidly extracted into fresh buffer on ice. A series of 2-mm coronal blocking cuts were rapidly made using a custom-designed marmoset brain matrix. Slabs were transferred to a dish with ice-cold buffer, and regions of interest were dissected using a marmoset atlas as reference. Samples of somatosensory and motor cortex were flash-frozen in RNAlater (Invitrogen) or immediately processed for cell dissociation.

#### RNA-seq of genetically-identified projection neuron populations

Pools of 5,000-10,000 cortical pyramidal neurons were sorted directly into TRIzol-LS buffer (Invitrogen) and RNA was extracted according to the manufacturer’s protocol. 10 ng total RNA was used for library preparation using the SMART-Seq v4 Ultra Low Input RNA Kit (Clontech) and the Nextera XT DNA Library Preparation Kit (Illumina) according to the manufacturer’s protocols. All libraries were sequenced according to the manufacturer’s protocols on the HiSeq 2500 system (Illumina), using 125bp paired-end reads, at a depth of >20 million reads per library.

#### ATAC-seq of genetically-identified projection neuron populations

Pools of 3,000-5,000 cortical pyramidal neurons were sorted into Opti-MEM media (Life Technologies). Nuclei were extracted and libraries prepared following a previously published protocol (Buenrostro et al., 2013). All libraries were sequenced according to the manufacturer’s protocols on the HiSeq 2500 system, using 50bp single-end reads, at a depth of ∼50 million reads per library.

#### WGBS of genetically-identified projection neuron populations

Pools of 5,000-10,000 cortical pyramidal neurons were sorted into PBS. The EZ DNA Methylation-Direct Kit (Zymo Research) was used to perform bisulfite conversion and libraries were prepared with the EpiGnome Methyl-Seq Kit (Illumina). All libraries were sequenced according to the manufacturer’s protocols on the HiSeq 2500 system, using 125bp paired-end reads, at a depth of >200 million reads per library.

#### Single-cell/single-nucleus RNA sequencing Mouse

Somatosensory and motor cortex from wild-type animals was dissected and dissociated at P1, P7, and P21. Tissue dissociation was performed as described (Arlotta et al., 2005), and live cells were isolated by FACS sorting as DAPI-negative, Vybrant DyeCycle Ruby (Thermo Fisher)-positive events. Libraries were prepared using the 10x Genomics Chromium Single Cell 3’ kit v2 (10x Genomics) according to the manufacturer’s protocol.

#### Marmoset

Somatosensory and motor cortex from wild-type animals was dissected and flash-frozen in RNAlater (Invitrogen). Nuclei were extracted by a previously-published protocol (Ma et al., 2018), and debris was removed by FACS purification of DAPI-positive nuclei. Nuclei were sequenced using the 10x Genomics Chromium Single Cell 3’ kit v2 (10x Genomics) according to the manufacturer’s suggested protocol for nuclei.

#### Single-cell/single-nucleus ATAC sequencing Mouse

Somatosensory and motor cortex from wild-type animals was dissected at P1, P7, and P21. Libraries were prepared as described in (Lafave et al., 2020), a modification of (Cusanovich et al., 2015). Briefly, cells were fixed with 0.1% formaldehyde and incubated at room temperature for 5 min. The fixation was stopped by adding glycine to the final concentration of 125 mM. The sample was incubated at room temperature for 5 min and washed in PBS. The cell concentration was counted, and approximately 1600-2000 cells per well were distributed into each well of a 96-well plate. Cells were transposed with 96 uniquely barcoded Tn5 at 37°C for 30 min with shaking at 300 rpm. The reaction was stopped by adding 0.5 M EDTA and incubated at 37°C for 15 min. All the cells were then pooled and MgCl2 was added to the pooled sample to quench EDTA. The sample was re-distributed onto another 96-well plate with 20 cells in each well by FACS sorting. Reverse crosslinking buffer and barcode PCR primers were added to each sample. The plate was incubated at 55°C for 16 hours for reverse crosslinking. Tween 20 was then added to quench SDS before PCR amplification.

The PCR reaction was carried out at the following conditions: 72°C for 5 min (extension), 98°C for 5 min, and then thermocycling at 98°C for 10 s, 70°C for 30 s and 72°C for 1 min for 12-15 cycles. Libraries were pooled and purified using Qiagen MinElute PCR purification column. The libraries were quantified using KAPA library quantification kit. Libraries were sequenced on the Next-seq platform (Illumina) using a 150-cycle kit (Read 1: 47 cycles, Index 1: 36 cycles, Index 2: 36 cycles, Read 2: 47 cycles).

#### Marmoset

Somatosensory and motor cortex from wild-type animals was dissected and dissociated with the Worthington Papain Dissociation System (Worthington Biochemical Corporation), and live cells were isolated by FACS sorting as DAPI-negative, Vybrant DyeCycle Ruby (Thermo Fisher)-positive events. Libraries were prepared using the 10x Genomics Chromium Single Cell ATAC kit (10x Genomics) according to the manufacturer’s protocol.

#### Immunohistochemistry

To confirm class specificity of labelling, CRE recombination was induced at E17.5 by tamoxifen administration, and mice were sacrificed at E18.5, P1, P3, P7, P21 and P48 for co-immunostaining with the canonical layer markers CUX1, CTIP2, and SATB2.

Brains were fixed and stained using standard methods (Arlotta et al., 2005). Briefly, mice were deeply anesthetized with tribromoethanol and perfused transcardially with phosphate-buffered saline (PBS) followed by 4% paraformaldehyde in PBS (PFA). Brains were post-fixed in 4% PFA overnight, washed in PBS, embedded in low-melting point agar, and sectioned at 20 micrometers using a Leica VT 1000s vibrating microtome.

Immunohistochemistry was performed as previously described (Arlotta et al., 2005). Primary antibodies and dilutions were as follows: mouse anti-Satb2, 1:50 (Abcam, ab51502); rat anti-Ctip2, 1:100 (Abcam, ab18465); rabbit anti-Cux1, 1:300 (Santa Cruz, sc-13024). Secondary antibodies were from the Molecular Probes Alexa series. Imaging was performed using a Nikon 90i fluorescence microscope equipped with a Retiga Exi camera (Q-IMAGING). Analysis was done with Volocity image analysis software v4.0.1 (Improvision).

#### Bioinformatics analysis

##### Data processing

###### Bulk RNA-Seq

Raw reads were trimmed using trimmomatic (Bolger et al., 2014) version 0.33, removing 8bp from the 5’ and 25 bp from the 3’ end. Subsequently, reads were aligned to the ENSEMBL NCBI37 (mm9) genome build (downloaded from the Illumina iGenomes file collection), using tophat2 (Kim et al., 2013) version 2.0.13 with default parameters. Subsequently, differential gene expression analysis and FPKM quantification was performed using cuffdiff (Trapnell et al., 2013) version 2.2.1 for all pairwise comparisons. Differentially expressed genes were defined as all genes that showed a significant (FDR≤0.05) change in gene expression with ≥1.5 log_2_ fold-change in at least one comparison, and were expressed at levels greater than 10 FPKM in at least one condition within that comparison, resulting in n=4,419 differentially expressed genes across the entire dataset.

Next, genes were clustered using k-means clustering on the log_2_ transformed and z-scored FPKM values of all differentially expressed genes using 100 random starts. In order to determine the number of clusters, we used the gap-statistics as implemented in the R package cluster (Maechler, 2019) in combination with the Tibshirani 2001 method (Tibshirani et al., 2001) based on the standard deviation evaluating k=2 to 20, identifying 12 clusters in total. Finally, we classified each cluster manually based on its expression dynamic into one of 5 categories: shared developmental clusters (1-5), Cux2 CPN-specific (6-8), Tle4 CThPN-specific (9-11), and other (12), as shown in Fig. 1C.

##### Bulk ATAC-Seq

ATAC-Seq raw reads were aligned to the genome build NCBI37 downloaded from the Illumina iGenomes collection using bowtie2 (Langmead and Salzberg, 2012) with default parameters. Subsequently, aligned reads were filtered for duplicates using MarkDuplicates from the picard software toolbox (broadinstitute.github.io/picard) (Broad Institute of MIT and Harvard, 2020). Next, we performed peak calling for each sample group using the IDR framework (Li et al., 2011) in combination with the macs2 peak caller, with two independent biological replicates in each group. All peaks detected at an IDR ≤0.1 in each group were retained for further analysis. We then performed differential peak enrichment analysis across all pairwise group comparisons using the diffBind package (Ross-Innes et al., 2012; Stark and Brown, 2011) in combination with DESeq2 (Love et al., 2014). To that end, we employed the DBA_SCORE_TMM_READS_EFFECTIVE score for normalization and subsequent differential enrichment analysis. We defined all peaks exhibiting significant (FDR≤0.01) differential enrichment above a log_2_ fold change of 1.5 and a minimum enrichment ≥1 TMM normalized reads in at least one condition as differential, resulting in n=66,784 differentially enriched ATAC-seq peaks across the entire dataset. Subsequently, we averaged over all replicates for each group and transformed the resulting TMM value to log_2_ space. Next, we conducted k-means clustering on z-scored and log2 transformed TMM values on all differentially enriched regions using 100 random starts. We again used the gap statistics in combination with the Tibshirani SE method to identified 13 clusters. We then classified each cluster as developmental, neuron class-specific, or other according to its dynamic enrichment patterns (Fig. 1D).

Finally, we associated each differentially active ATAC-Seq peak with the closest ENSEMBL gene TSS using the ChIPpeakAnno package (Zhu et al., 2010). Selected gene names based on this association are shown in Fig. 1D.

##### Bulk WGBS

Raw sequencing reads were trimmed 8bp from the 5’ end and 40 to 60bp from the 3’ end, depending on library quality. Next, reads were aligned to the genome build NCBI37 using bsmap (Xi and Li, 2009) version 2.9 with parameter settings -v 0.1 -s 16 -q 20 -w 100 -S 1 -u –R. Aligned data was then filtered for PCR duplicates using the MarkDuplicates function implemented in the picard toolbox. Next, CpG methylation calling was performed on the duplicate filtered data using the mcall function implemented in the MOABS suite (Sun et al., 2014) with default parameters. We identified n=44,127 differentially methylated regions, using p-value ≤ 0.01 and methylation difference ≥ 0.3. Clustering analysis identified 10 clusters with distinct temporal and cellular context dynamics.

##### Single-cell ATAC

Base calls were converted to FASTQ format using bcl2fastq (Illumina). Raw sequencing reads were trimmed using custom python scripts to remove adapter sequences. The reads were aligned to mm10 or CalJac3 genome using Bowtie2 (Langmead and Salzberg, 2012) with maximum fragment length set to 2 kb, and all other default settings (bowtie2 -X2000 --rg-id). The data were demultiplexed tolerating one mismatched base within barcodes. Mitochondrial, unpired and low-quality reads were removed using SAMtools (Li et al., 2009) (samtools view –b -q 30 -f 0x2). Duplicate sequences were removed using the picard toolkit (Broad Institute of MIT and Harvard, 2020).

#### Data analysis

##### Identification of differentially methylated regions (DMRs)

DMRs were identified using the R package DSS (Park and Wu, 2016). To that end, we performed all pairwise comparisons across sample groups. For each of these pairwise comparisons, we applied the following 3 functions from the DSS package to the appropriate biological replicates. First, utilized dmlTest with smoothing=T and smoothing.span=200. Next, we identified differentially methylated CpGs using the callDML function with a threshold of p=0.001. Finally, we identified DMRs using the callDMR function directly using the posterior probability that the methylation difference exceeds a certain value. We set the parameters delta=0.3, p.threshold=0.01, minCG=3 and dis.merge=400; all other parameters were left at their default settings. Next, we merged all DMRs identified across the pairwise comparisons into one DMR set, collapsing DMRs that overlap by at least one base pair into a single DMR. For further downstream analysis and visualization, we employed the methylKit package (Akalin et al., 2012). In particular, we computed the methylation level and coverage of each DMR in each sample, defined as the weighted average of CpG methylation levels weighted by coverage. We then only retained those DMRs that were covered by more than 5 reads in at least 3 samples. Next, we averaged the methylation levels of each DMR across replicates and assigned the DMRs to different groups based on their methylation differences between the replicate averaged, DMR-level methylation values. DMRs that exhibited an absolute methylation difference ≥ 0.3 between any pair of samples and exceed a size of 100bp were defined as differentiation DMRs.

We then performed clustering using k-means, initially identifying 12 clusters that we collapsed upon further inspection into 10 distinct clusters. We again annotated each cluster according to its dynamic enrichment patterns (Fig. 1E).

We determined global CpG and non-CpG methylation levels as the fraction of methylated CpGs over the total number of detected CpGs (and correspondingly for non-CpGs) for all ATAC-Seq regions overlapping with DMRs and covered by at least 10 reads in 80% of the samples using the function in the regionCounts function in methylKit package, and report the corresponding feature methylation values in Fig. S4C.

#### Multidimensional scaling analysis

We performed multidimensional scaling using the cmdscale R function with 1 minus the absolute Person Correlation Coefficient as metric, reducing the dimensionality to 2 dimensions. We then computed the distances of individual samples shown in Fig. 1G and corresponding text as the 2-dimensional Euclidean distance. As input, we used the log2+1 transformed FPKM values of all expressed genes (≥10FPKM in at least one condition) (RNA-Seq), the log2+1 and quantile normalized ATAC-Seq TMM values (ATAC-Seq) and the methylation level of all 1kb tiles of the mouse genome covered by at least five reads in more than 2 samples. In order to estimate confidence intervals for the computed Euclidean distances reported in the text, we randomly removed 20% of all genes/peaks/tiles, repeated the multidimensional scaling and recomputed the distances. We repeated this process 1000 times for each assay and report the corresponding confidence level estimates in the text.

#### Specificity analysis for bulk RNA-Seq clusters

We report specificity analysis for differentially expressed TFs as well as gene expression clusters. These specificity analyses are conducted using 3 distinct datasets that were processed in the following manner:

*FANTOM5* (*19)*: We computed the expression specificity of all genes and all TFs in each expression cluster across 294 mouse cell types and tissues based on CAGE data from the FANTOM5 consortium (FANTOM Consortium and the RIKEN PMI and CLST (DGT) et al., 2014). To that end, we downloaded the CAGE-tag data for promoter regions from the FANTOM5 cell and tissue collection from http://fantom.gsc.riken.jp/5/. We then collapsed all CAGE-tag peaks for each gene by summing up the tag counts, including only primary cell types. Subsequently, we collapsed replicates for each cell type or tissue by averaging.

*Allen Brain in situ* (Lein et al., 2007; Thompson et al., 2014): The Allen brain data was downloaded from their website (http://www.alleninstitute.org).We obtained *in situ* hybridization counts for the developing mouse brain at 7 distinct fetal time points and 11 different brain substructures through direct correspondence with http://www.alleninstitute.org. We then intersected the resulting list of genes with the list of transcription factors/genes in each expression cluster and determined the expression specificity of each TF/gene across the 77 conditions (see below for further details) from this atlas. We plot the distribution of specificities for each cluster in Fig. 2C.

*Mouse single cell RNA-Seq data*: Here, we used the log2 normalized expression values averaged over the all cells in a particular cell cluster using the AverageExpression() function in Seurat. We then utilized these pseudo-bulk expression values for each gene across all identified cell types as input for the specificity analysis.

Similarly, we computed the expression specificity of all genes and all TFs in each cluster across 294 mouse cell types and tissues based on CAGE data from the FANTOM5 consortium (FANTOM Consortium and the RIKEN PMI and CLST (DGT) et al., 2014). To that end, we downloaded the CAGE-tag data for promoter regions from the FANTOM5 cell and tissue collection from http://fantom.gsc.riken.jp/5/. We then collapsed all CAGE-tag peaks for each gene by summing up the tag counts.

We then computed expression specificity for each TF/gene following previous approaches using the tau specificity measure (Ravasi et al., 2010) according to (Kryuchkova-Mostacci and Robinson-Rechavi, 2017):

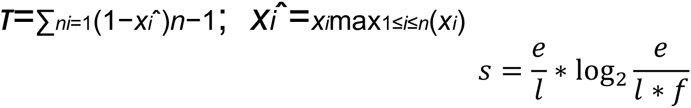

With e being the vector of gene expression values of length f and l being *l* = ∑*e_i_*.

#### Specificity analysis for single-cell/single-nucleus ATAC clusters

In order to assess the specificity of each ATAC peak in the bulk or single cell/nucleus ATAC dataset, we downloaded pre-computed ATAC peak specificity scores computed over more than 80 distinct cell types of an entire mouse using single cell ATAC-seq data from Cusanovich et al. (Cusanovich et al., 2018).

We then intersected our peak library with the Cusanovich dataset and report the Cusanovich specificity values for all peaks that overlap with at least one peak in the Cusanovich dataset.

##### Bulk ATAC-peak overlap with DHS data

In order to create a catalog of gene regulatory elements in the mouse genome, we downloaded a set of DNAse HS I peak tracks from the mouse ENCODE consortium (Yue et al., 2014). Subsequently, we collapsed replicates for each condition and required that each peak was present in at least two replicates. This step resulted in DNAse I tracks for 35 distinct primary mouse cell types and tissues. Next, we merged all DNAse I tracks into a union peak set using the reduce() function in the IRanges R package (Lawrence et al., 2013). Subsequently, we size standardized the resulting union peak set to 350bp by extending 175bp from the center of each peak. Next, we assigned a binary value for each peak in each of the 35 cell types, depending on whether or not the peak was present in the individual cell type-level peak set. We then used this union peak set and overlapped all ATAC-peaks from the mouse bulk ATAC-seq dataset with this library. We report the percent of peaks in each cluster that overlap DNAse I HS sites in each cell type.

##### Definition of genomic features

CpG islands were defined as previously described (Bock et al., 2012). Promoters were defined as all NCBIm37 ENSEMBL version 67 TSS, extended by 1kb upstream and downstream.

##### Transcription factor binding site (TFBS) density analysis

TFBS density was determined based on FIMO motif instance matches below or equal to a p-value of 10^−4^. The density was then calculated as the total number of motif instances across all member regions of the cluster divided by the size standardized peak length in bp. The distribution of TFBS densities across all motifs is then reported for each ATAC-Seq peak cluster.

#### Entropy analysis

In order to compute the Shannon Entropy of the size standardized ATAC-Seq peak sequence we used the *entropy()* function in the sequtils (Cock et al., 2009) python package.

#### Integrative analysis

In order to evaluate the concordance of changes in the transcriptome, open chromatin and DNA methylation landscape, we associated each ATAC-peak or DMR with its nearest gene within 100kb upstream or downstream. Peaks/DMRs without any assigned gene were not considered. Subsequently, we performed a hypergeometric test between all pairs of ATAC-peak/DMR clusters and bulk expression clusters in gene space to assess the significance of overlap. Following multiple-testing correction using the Benjamini-Hochberg Method (Benjamini and Hochberg, 1995), we report the odds-ratio of associations significant below a q-value of 0.001 in Fig. S6C-D.

#### Phylogenetic conservation analysis

We performed phylogenetic conservation analysis for ATAC-seq cluster groups by computing the average placental mammal phyloP scores (Pollard et al., 2010) for each region. We then plot the distribution of these mean scores.

#### Signature gene set analysis

First, we associated each of our consensus DMRs with the nearest mouse ENSEMBL TSS within 100kb using the ChIPPeakAnno R package (Zhu, 2013; Zhu et al., 2010). Subsequently, we performed gene-set-level analysis and determined the mean methylation level of all DMRs associated with a member gene of each signature gene set using the aforementioned DMR-gene associations. We used signature gene sets for CThPNs, CPNs and sub-cerebral projection neurons (SCPNs) obtained from the DeCoN resource (Molyneaux et al., 2015) as well as manually curated gene sets for glial and interneuron cell types from published transcriptomic data (Cahoy et al., 2008; Zhang et al., 2014). We then report the mean methylation level of all DMRs associated with each gene set in Fig. S5C for each time point and cell type.

#### Global methylation level analysis

Global CpG methylation level was defined as the total number of detected methylated Cs in CpG context over the total number of CpGs sequenced. Similarly, we defined the global non-CpG methylation level comprising all other dinucleotide contexts.

#### Mouse single cell RNA-seq analysis

Mouse single cell RNA-seq data was processed using Cell Ranger version 3.0.1 (10x Genomics) using standard parameters and genome assembly GRCm38 downloaded from the 10X Genomics homepage. Following initial alignment and processing by cellranger count, all replicates across all time points were aggregated using the cellranger aggr function, downsampling the individual libraries to a similar overall coverage by cell.

All following analyses were conducted in R using the Seurat package (version 2.3.4) (Butler et al., 2018; Stuart et al., 2019). Only cells with at least 1000 genes, a mitochondrial read fraction below 10% and not more than 6000 genes or 15000 UMIs were retained.

Subsequently, we initially performed cell clustering and subtype identification separately for each time point P1, P7 and P21. For that purpose, we subset the data for each time point and then apply the following workflow: Expression data was normalized using the LogNormalize method and variable genes were identified based on the mean/dispersion relation (initially using the following parameters x.low.cutoff = 0.05, x.high.cutoff = 3, y.cutoff = 0.05).

Next, the data was scaled using the ScaleData function, regressing out percentage of mitochondrial reads and UMI count per cell. We then compute the first 100 PCAs and examine the resulting Elbow Plot for variance explained by each PC. Based on that, we selected the first 35-40 PCs for subsequent dimensionality reduction by UMap (min_dist = 0.2 to 0.5) and cluster identification using the Seurat FindClusters (resolution = 0.4 to 0.6, nn.eps = 0.5) function.

We conducted several rounds of PCA, clustering and Umap embedding, successively removing clear outlier cell clusters based on the UMAP (4 for P1, 2 for P7 and 2 for P21).

Following cleanup and cluster identification by time point, we analyzed all time points jointly, assigning the cell original cluster definitions based on the individual time point analysis. For that purpose, we used as cutoffs in the variable gene feature analysis: x.low.cutoff = 0.025, x.high.cutoff = 3, y.cutoff = 0.05, 50PCs, a resolution of 0.9 in the clustering analysis and a min_dist=0.4 in the umap.

Next, we performed cell type identification for each cluster using a set of manually curated marker genes for each cell type. In order to assign identities, we examined the expression of all marker genes individually using the UMAP. In addition, we computed the AverageExpression for each cluster and examined the pseudo-bulk profiles. In particular, we computed joint cell type scores for each cluster and potential identity by normalizing the average cell type scores for each cluster to the maximum observed score for each cell type separately. Moreover, we performed hierarchical clustering, correlation analysis, PCA and MDS evaluation of the pseudo-bulk profiles to identify outlier clusters and investigate the relationship between cell clusters in more detail. Based on these analyses, we were able to assign specific identities to most cell clusters. However, several clusters appeared to have mixed identities (in particular clusters of interneurons). We thus conducted several rounds of subclustering for these to refine the identities of these cells. We collapsed distinct clusters with the same subtype identity (e.g. cluster CPN_1 and CPN_4 etc.) into a single subtype cluster to reduce the complexity for subsequent analysis (final cluster N=32).

#### Differential expression analysis

In order to identify differentially expressed between time points within distinct cell type classes, assigned each cell cluster to 7 sets of general cell classes that we analyzed separately: excitatory neurons, inhibitory neurons, astrocytes, oligodendrocytes, neurons, glia, and all cell types.

We then performed differential expression analysis in pairwise fashion between P1 and P7, P1 and P21, and P7 and P21 for each of the aforementioned cell type classes using the *FindMarker* function in Seurat and the MAST method for differential expression testing.

In this manner, we obtained 7 distinct lists of genes differentially expressed across development. Next, we averaged the expression of all cells within one cluster using the AverageExpression function in Seurat. In order to avoid biases driven by cluster complexity and cell number, we downsampled each cluster to a maximum of 500 cells per cluster prior to averaging. Next, we performed clustering on z-score transformed expression values of each of the 7 differentially expressed gene sets separately. For that purpose, we used the clusGap function in the R package cluster (Maechler, 2019) using kmeans with a maximum of 20 clusters and the B parameter set to 60. We then select the final number of clusters based on the Tibshirani 2001 criterion (Tibshirani et al., 2001) for the standard deviation as implemented in the cluster package. Finally, we perform kmeans clustering with 100 random starts using the identified number of clusters and average over the z-scored genes in each cluster.

Lastly, we performed expression specificity analysis for all gene clusters in a similar manner as for the bulk data, using the Allen Brain in situ dataset, the FANTOM5, and (for marmoset) the mouse scRNA-Seq dataset for comparison for all differentially expressed genes as well as for transcription factors only.

#### Marmoset single nucleus RNA-seq analysis

Marmoset snRNA-seq data was processed using the Cell Ranger version 2.1.0 and aligned to a pre-mRNA custom build transcriptome of assembly version ASM275486v1.93. We selected this assembly due to the improved transcript and gene annotation available compared to previous assemblies. All replicates were first processed independently using Cell Ranger count and then aggregated, downsampling all libraries to the same complexity per cell. Subsequently, the data was processed using Seurat following a similar workflow as for the mouse scRNA-seq, retaining only cells with more than 1000 genes and genes detected in more than 10 cells. Initially, variable genes were again identified using the mean/dispersion relation with the following parameters: x.low.cutoff = 0.05, x.high.cutoff = 3, y.cutoff = 0.05. During initial QC of the 5 individual marmoset libraries, we observed a separation by experimental batch, where all three libraries (D0 and Y2) from experimental batch 1 and the two libraries from batch 2 (D0 and Y2) grouped together. Given that developmental time point and experimental batch are not confounded in our experimental design, we performed batch correction using CCA as implemented in the Seurat package. Here, we discarded all cells where the variance explained by CCA is <2-fold. Following correction, a clear separation by time point and cell type became apparent, as originally observed when processing each experimental batch separately. We thus proceeded with the CCA corrected data following the same strategy as for the mouse scRNA-Seq data, performing PCA and using 20 dimensions in subsequent analyses, identifying clusters with a resolution of 1.8 and creating a UMAP embedding with a min_dist of 0.3.

Subsequently, data analysis was conducted similar to the mouse scRNA-Seq data, assigning cell types and cell classes, performing differential expression analysis between time points within each of 7 cell classes, collapsing clusters with similar subtype identity and performing gene clustering within each cell group to identify gene clusters.

#### Read alignment and pre-processing

Base calls were converted to FASTQ format using bcl2fastq (Illumina). Raw sequencing reads were trimmed using custom python scripts to remove adapter sequences. The reads were aligned to mm10 or CalJac3 genome using Bowtie2 (Langmead and Salzberg, 2012) with maximum fragment length set to 2 kb, and all other default settings (bowtie2 -X2000 --rg-id). The data were demultiplexed tolerating one mismatched base within barcodes. Mitochondrial, unpaired and low-quality reads were removed using SAMtools (Li et al., 2009) (samtools view – b -q 30 -f 0x2). Duplicate sequences were removed using the picard toolkit (Broad Institute of MIT and Harvard, 2020). In order to counteract differential complexity across the libraries, each library was sampled to a similar fragment depth per cell.

#### Single-cell and single-nucleus ATAC analysis

Mouse and marmoset libraries were first downsampled to similar overall complexity in terms of reads/cell across all conditions. Cleaned data was processed with the scasat (Baker et al., 2019) pipeline using macs2 as peak caller with parameters set to -q 0.2 --nomodel – nolambda, giving rise to a peak − cell matrix count matrix. For genome size, we used -g mm for mouse and -g 2.1e+9 for marmoset. We then size-standardized all peaks to 500 bp and recomputed the peak − cell count matrix, only considering reads overlapping the size-reduced peaks.

Only cells with at least 1,000 reads and 300 peaks from the master list overlapping with at least one read, but not more than 15,000 reads and 8,000 peaks were retained for analysis. In addition, only peaks present in at least 20 cells were retained.

Subsequently, this matrix was processed using R implementing a custom processing pipeline based on the strategy outlined in Cusanovich et al. 2018 (Cusanovich et al., 2018) and refined by Andrew Hill (Hill, 2019), following the log-LSI workflow. Briefly, the count matrix is first binarized and then transformed using the TF-IDF method (Cusanovich et al., 2015) log scaling the results. Next, PCA is performed on the transformed matrix using 50 dimensions, following cell cluster identification based on Seurat’s *FindCluster()* function with resolution set to 0.5 and UMAP embedding with a min_dist of 0.3. Importantly, we split each cluster containing cells from different time points into separate clusters for each time point and then filter out all clusters with less than 100 cells.

#### Cell type identification

In order to reliably identify distinct cell types, we next computed gene activity scores for all genes based on the presence of ATAC peaks. To that end, we utilized the R package cicero (Pliner et al., 2018) since it implements an approach not only considering peaks located within the promoter/gene body of each gene, but also weights the contribution of each peak to the overall gene score based on the correlation of the gene peaks with each other. To that end, we imported the preprocessed data into a cicero atac_cds object providing the cluster id and UMAP coordinates defined based on the aforementioned LSI analysis. Next, we estimated library size factors and reliable peaks using default parameters followed by running cicero’s main function with default parameters to compute the connectivity graph of all peaks. Next, we annotated all peaks with the *annotate_cds_by_site* function using transcription start coordinates for all ensemble genes retrieved from biomaRt defining the region +/− 5kb of each TSS as promoter region. With this annotation in place, we constructed the raw gene activity matrix using the build_gene_activity_matrix function, followed by normalizing the resulting scores using the function *normalize_gene_activities*.

Next, we average the computed gene scores for each cell cluster and normalize the aggregated score for each gene to the maximum across all pseudo-bulk cell clusters. We then again compute cell type scores for each scATAC cell cluster by averaging a set of known marker genes (same as for scRNA-Seq) for each cluster and plotting results along with hierarchical clustering, PCA and MDS analysis of the pseudo-bulk gene scores. Based on these analyses, we remove outlier clusters (e.g. very low complexity) and assign a final cell type annotation. Based on this cleaned dataset, we perform differential accessibility analysis, aggregating the signal of all cells in each cluster using cicero’s *aggregate_by_cell_bin* function followed by a negative binomial based differential accessibility test implemented in *differentialGeneTest* function using the cluster id as factor to test. In addition to the cell-cluster based test, we also perform a second test on the time point variable to identify specifically those peaks variable across distinct time points. We then define all peaks significant below a q-value of 0.5 in either of the two analysis as dynamic peak set for subsequent analysis. The results are not particularly sensitive to this threshold, as we tested qvalue thresholds between 0.1 and 0.5. However, given the limited power for each peak, we decided to include all peaks with evidence for differential accessibility. We opted to an 0.5 threshold as this gave us a similar fraction of variable peaks as the bulk ATAC analysis conducted earlier.

All subsequent analyses were conducted on this dynamic peak set, after excluding all cells of unidentified identity. Next, we normalized the read counts for each cell by the respective size factor and averaged the resulting values across all cells within one cluster, giving rise to pseudo-bulk profiles. Next, we normalized these profiles for each peak across all pseudo-bulk clusters by dividing each row (corresponding to peaks) by the 95^th^ quantile across all cell clusters, capping all values at 1. This gave rise to a normalized accessibility score in the unit interval. These values were then used for clustering analysis, following the same strategy as for the scRNA-seq, again assigning each cell cluster to one of 7 categories and then performing k-means with 100 random starts and clustering on the respective subset of cell clusters for all peaks in the dynamic peak set. Again, the cluster number was determined using the clusGap package and Tibshirani 2001 SE criterion. The resulting peak clusters were then subjected to the same characterization as the peak clusters from the bulk scATAC analysis. For the specificity analysis, the cluster coordinates were lifted from mm10 to mm9 using the USCS liftover tool (http://genome.ucsc.edu/) (Kent et al., 2002).

#### Defining early, late, and cell type-specific clusters

In order to assign clusters to a particular activity pattern to individual region clusters, we predefined a set of patterns according to possible dynamics of interest: These included temporal (e.g. specific to P1, P7, P21, or P1_P7 or Y0 and Y2, excitatory neurons, inhibitory neurons as well as specific to a particular cluster). These patterns were summarized in a prototype binary indicator matrix of dimensions number of cell clusters times number of patterns. For each pattern and cell cluster, the indicator matrix cell was set to 1 if that cell cluster was associated with that pattern. For example, setting the indicator variable for P1_CPN_1 and P1_CPN_3 both to 1 in the P1 pattern definition.

We then computed the cosine similarity between each of these pattern vectors with the normalized accessibility score vector for each region cluster, taking the average of accessibility score for all regions within one region cluster for each cell cluster separately. This analysis gave rise to a similarity measure of each cluster’s dynamic to a predefined set of temporal/cell type dynamics. Based on the maximum observed similarity to a predefined pattern, we then assigned the pattern label to the corresponding cluster.

#### Mouse single cell ATAC peak methylation analysis

Prior to DNA methylation analysis, we performed a liftover of the size-standardized (500bp) scATAC peaks from mm10 to mm9 using the UCSC liftover tool. Next, we computed DNAme levels in a similar manner as for the bulk data, using the scATAC peaks instead of the DMR regions as input, only considering regions that were covered by at least 5 reads in at least 8 of the 12 WGBS samples. We report the distribution of methylation levels for each of the scATAC region clusters across CPN and CThPN neurons at P1, P21 and P48.

#### TF binding site enrichment analysis

For each set of regions of interest (DMRs, ATAC peaks) we performed motif detection analysis using FIMO (Grant et al., 2011) with a p-value filter of less than 10^−4^ and a joint motif data base comprising the TRANSFAC Professional library (version 2011) (Fogel et al., 2005), and a set of previously published motifs by Jolma et al. (Jolma et al., 2013). All genomic regions were size standardized (if not already) prior to motif analysis to 500bp (scATAC, bulk ATAC) or 350bp for DNAse and DMRs.

In order to perform motif enrichment analysis, a suitable class of genomic background regions is needed. Since a large fraction of DMRs likely correspond to gene regulatory elements and overlap with DNAse I HS sites, we used the union of DNAse I HS sites across a large panel of mouse cell types and tissues from the mouse ENCODE project (Yue et al., 2014). To that end, we downloaded all DNAse I HS peak sets from GEO, size-standardized them to 350 bp, and created a union peak set using the GenomicRanges R package (Lawrence et al., 2013). Subsequently, we size-standardized the union peak set and performed motif detection analysis on the union set as described above. With this set of background regions in hand, we performed motif enrichment analysis for each DMR and ATAC-Seq peak cluster separately. Since motif enrichment analysis is fairly sensitive to differences in GC content, we performed the enrichment analysis in a GC content-stratified fashion. To that end, we binned all regions within one cluster into 20 bins in 0.05 intervals. Next, we determined the number of motif instances for each motif in our library within all regions in one GC content bin. We then performed an identical analysis for a set of randomly sampled DNAse HS I regions with a similar GC content distribution from the mouse ENCODE background region set, and repeated this process 500 times. We then summed across all GC content bins and computed an empirical p-value and enrichment level for each motif in each cluster. We retained all motifs enriched at a p-value below 0.01 and an enrichment over background level above 1.5. Next, we associated each motif with its corresponding transcription factors and retained for each transcription factor the motif with the highest enrichment over background ratio. Finally, we filtered the resulting list for those transcription factors that also show differential expression in at least one condition within our dataset, based on the criteria outlined in the gene expression analysis section.

#### Similarity in cell types across ages

To compute the similarities among cell types within each age group and compare them, we first merged single cell data into pseudo-bulk by computing the average expression for each cell type at each time point (using the AverageExpression function from Seurat package v3.1.0 in R v3.6.3). We then performed the principle component analysis (using prcomp function included in R as its base package), taking the top 10 principle components (ordered by the fraction of total variance explained), to project the data onto the 2-D using the UMAP algorithm (umap package v0.2.4.0 in R).

#### Gene ontology enrichment analysis

We prepared input gene lists for each bulk RNA-seq shared early and shared late cluster based on our gene clustering. Enrichment analysis was done using the Panther overrepresentation test available through the Gene Ontology Consortium (Ashburner et al., 2000; Mi et al., 2019; The Gene Ontology Consortium, 2019) web interface. Test type was set to Fisher’s exact test, and false discovery rate was computed for each term. Selected GO terms for the shared-developmental clusters are presented in Table S1-S2.

### Supplementary Text

It has been reported that the Cux2-CreERT2 line used here also labels a subset of cortical interneurons (Franco et al., 2012; Gil-Sanz et al., 2015). To evaluate the effects of this on our analysis, we performed single-cell sequencing of a total of 14,792 cells for the Cux2 CPN and Tle4 CThPN populations at three ages, labelled using the same induction strategy used for our primary analysis (Fig. S2). Of these, 239 cells were positive for expression of the interneuron marker Gad1 (1.61%), 339 cells were positive for the interneuron marker Gad2 (2.29%), and 118 cells were positive for both (0.798%), for a total of <5% cells positive for either marker. We conclude that the effect of interneuron contamination on our analysis is minimal.

---

**Fig. S1.**
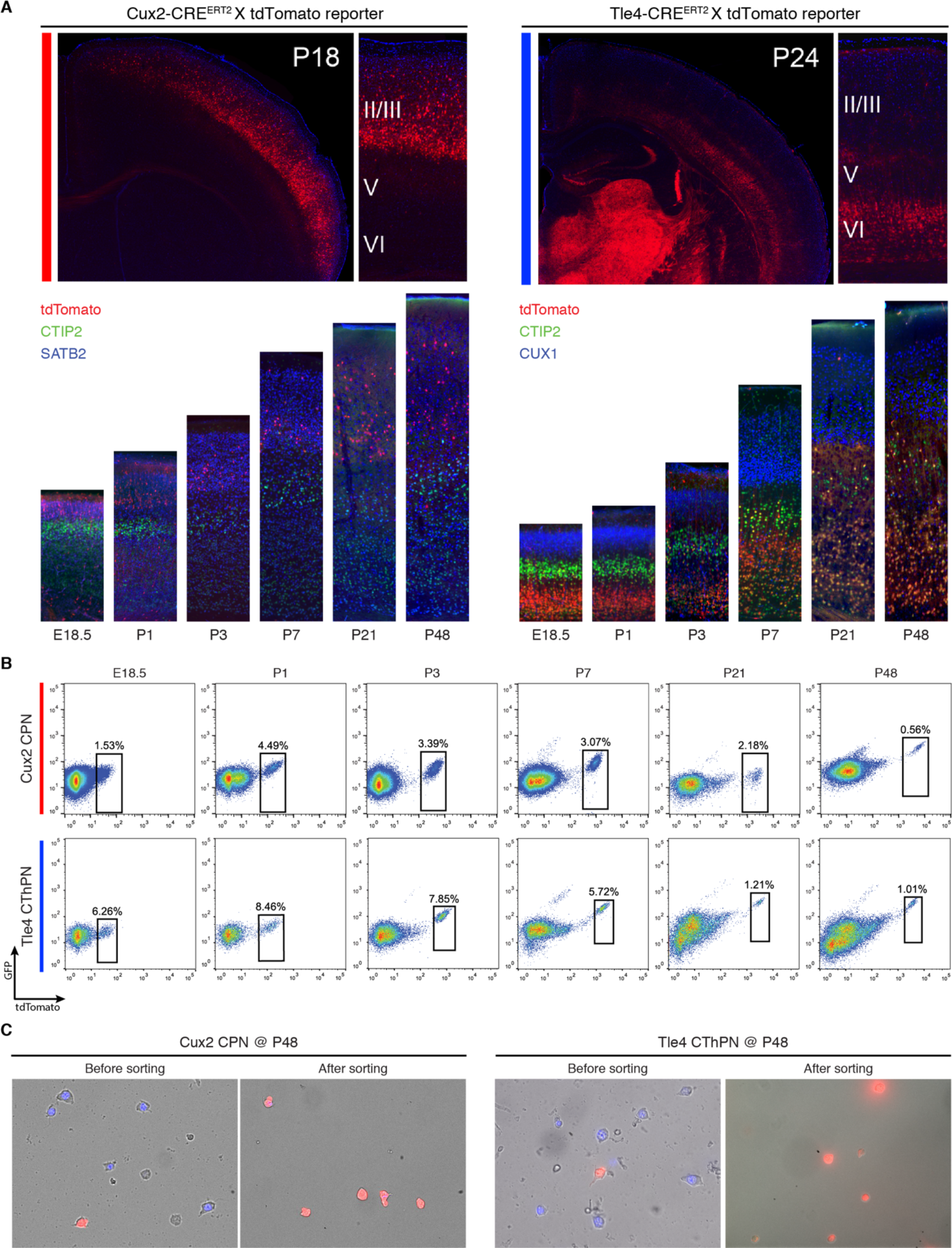
Isolation of genetically labelled CPNs and CthPNs. (**A**) Representative coronal sections showing correct laminar location of tdTomato^+^ cells in the somatosensory cortex at the different developmental stages. (**B**) Representative FACS results for purification of Cux2 CPN and Tle4 CthPN populations at the different developmental stages; boxes indicate collected populations. (**C)** Representative images of pre- and post-sorting Tle4 and Cux2 neuronal populations at P48, showing good purity and viability even at this older age.

**Fig. S2.**
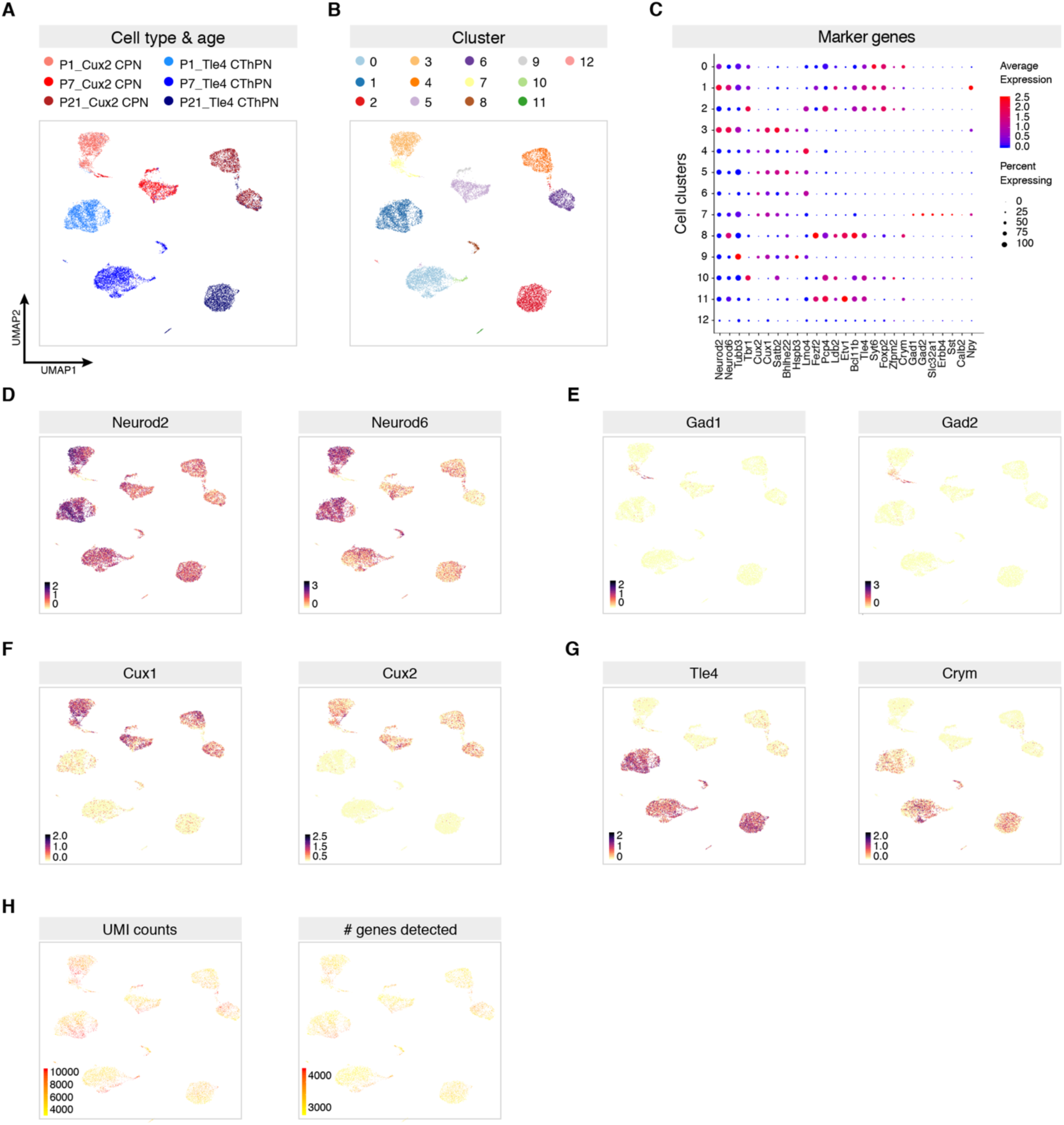
Verification of subtype identity of genetically-labelled mouse PN subtypes by single-cell RNAseq of FACS-purified Cux2 CPN and Tle4 CThPN. (**A**) UMAP showing distribution of cells by line and age. (**B**) UMAP showing cell clusters after k-means clustering. (**C**) Average expression of selected marker genes in each cluster. (**D-F**) UMAPs showing expression of selected marker genes in each cell, including pan-neuronal markers (D) and markers for interneurons (E), CPN (F), and CThPN (G). The majority of cells express markers corresponding to the expected pyramidal neuron subtypes, with only minor contribution of interneurons. (**H**) UMAPs showing number of UMIs and number of genes detected in each cell.

**Fig. S3.**
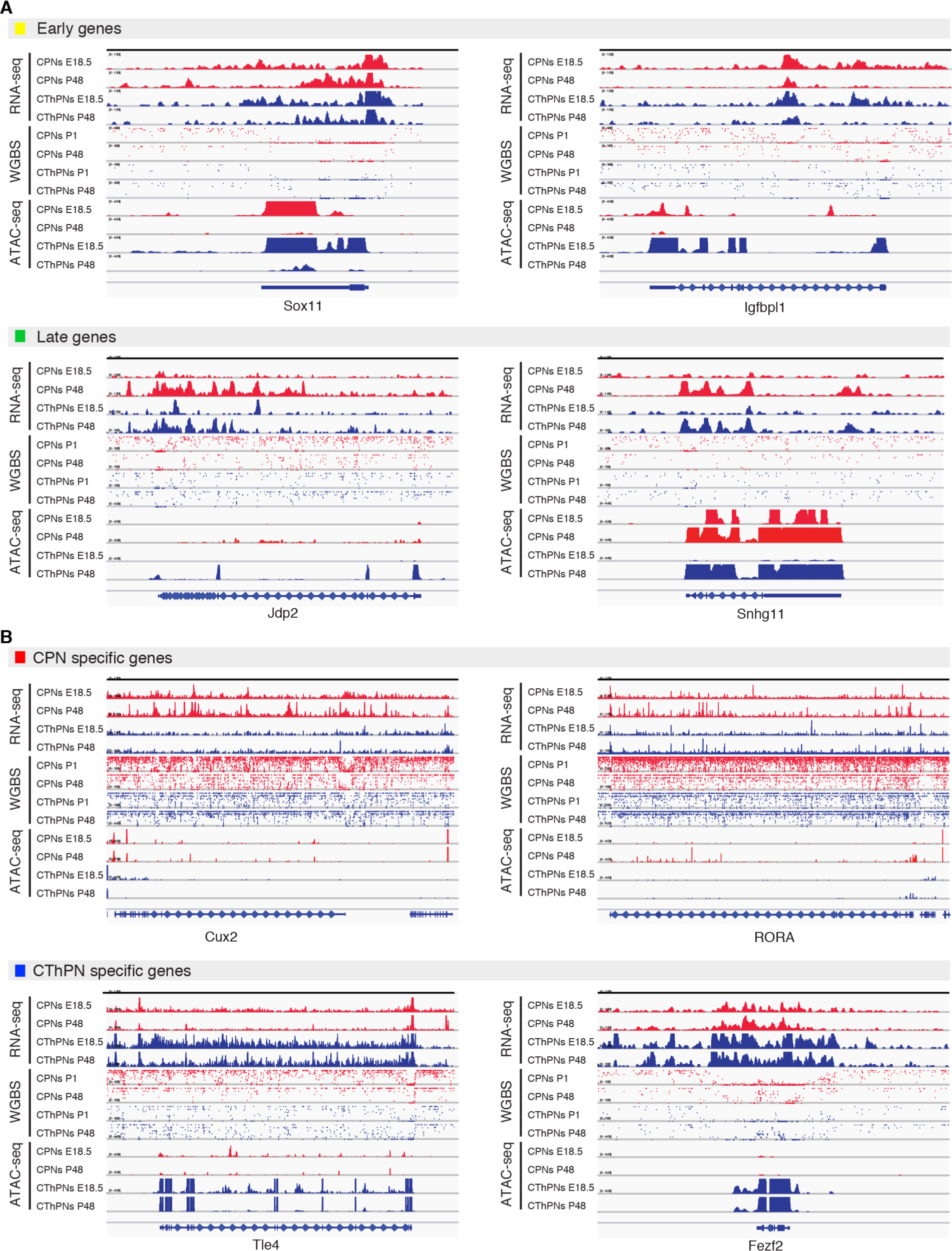
Example gene tracks for genes classified into different transcriptional categories. Examples showing Genome Browser views of RNA sequencing, ATAC-seq, and WGBS tracks, for examples of genes in (**A**) shared-developmental and (**B**) class-specific categories. Red: Cux2 CPNs; Blue: Tle4 CThPNs.

**Fig. S4.**
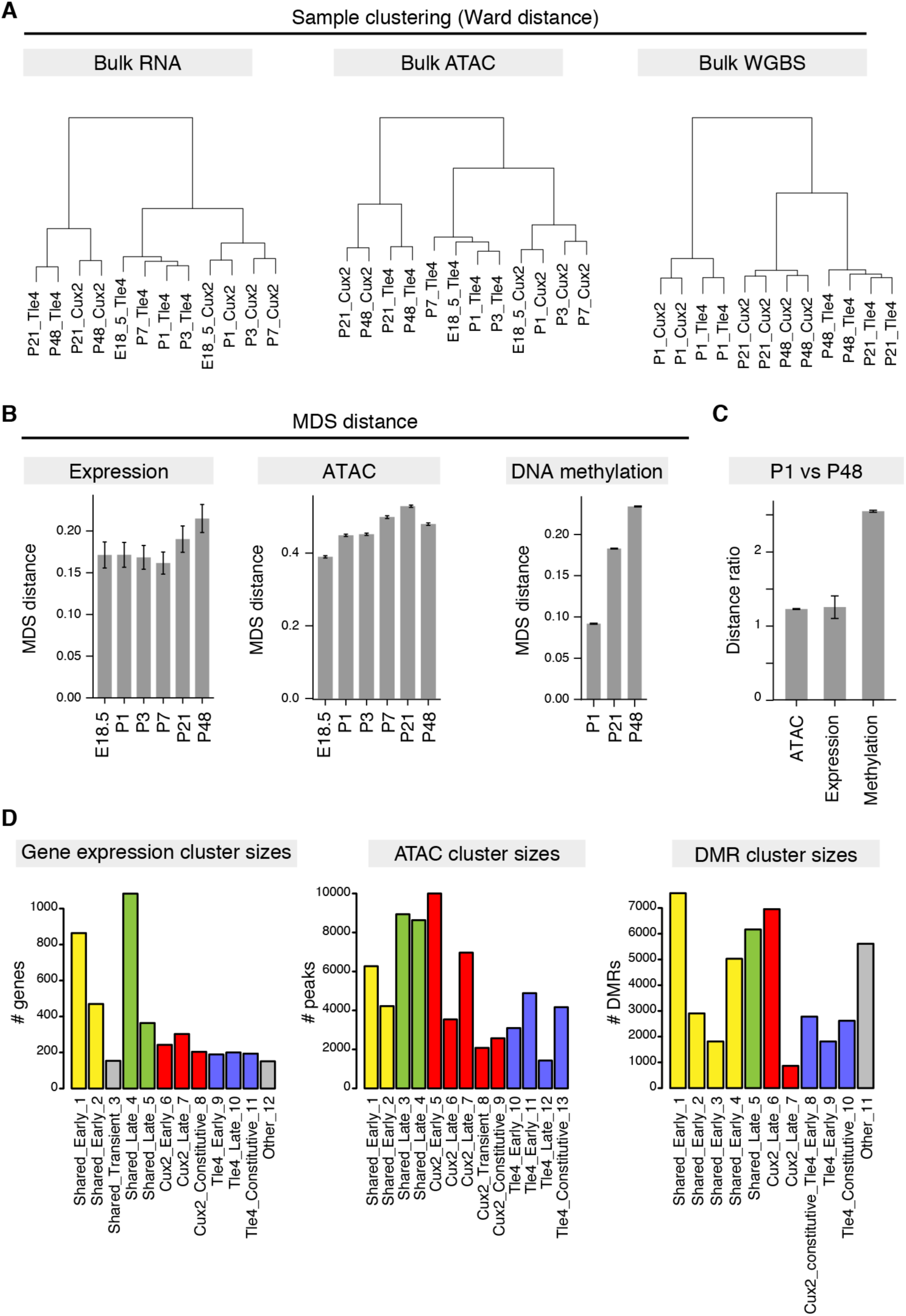
Characterizing bulk datasets. (**A**) Dendrograms showing relationships between samples from different ages for each dataset. (**B**) MDS distance values for the MDS analysis presented in Fig. 1G. (**C**) Ratio of distances at P1 vs. P48 for each dataset. (**D**) Number of features (genes, ATAC peaks, or DMRs) in each cluster for each dataset.

**Fig. S5.**
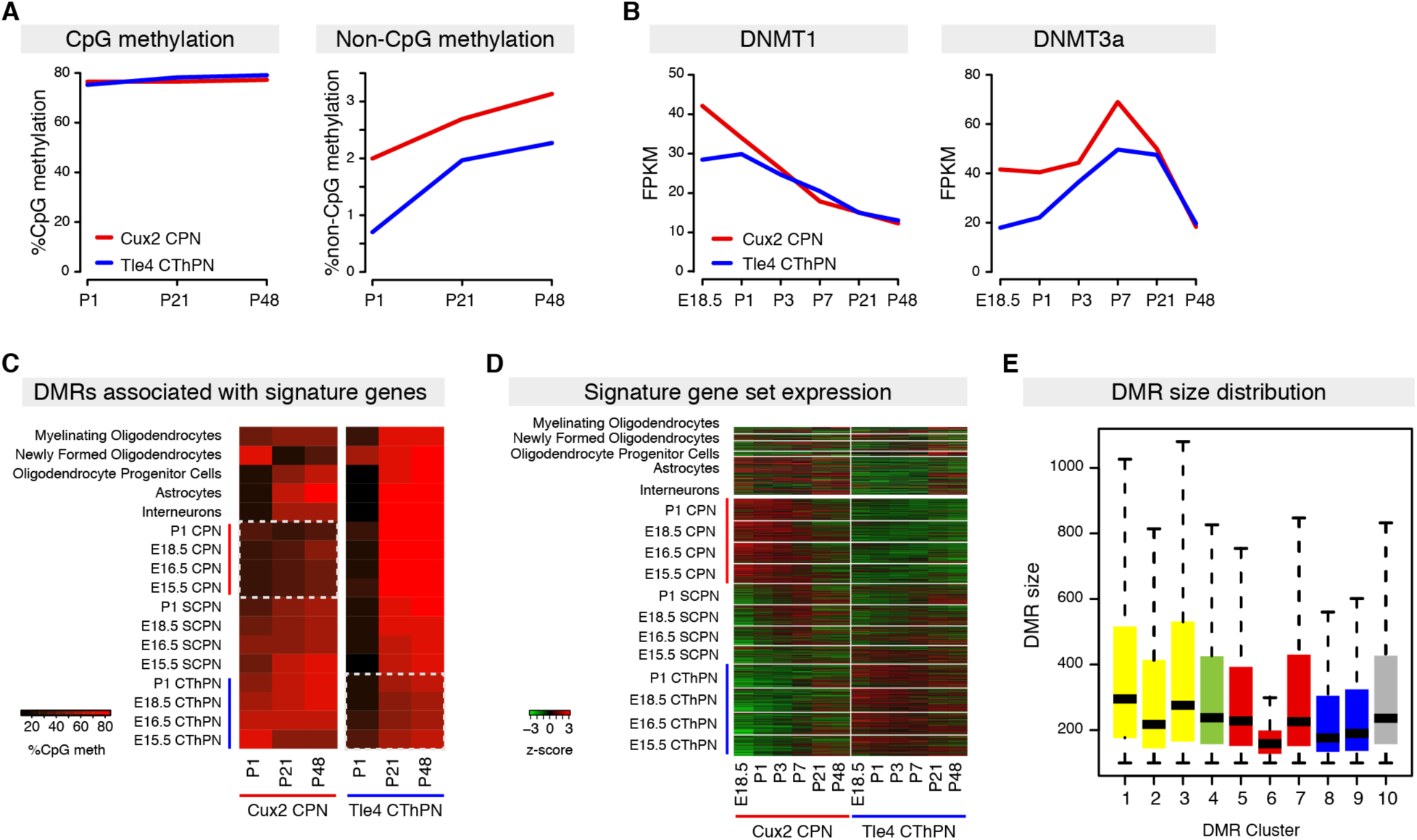
DNA methylation at dynamic loci. (**A**) Global DNA methylation levels at CpG (left) and non-CpG sites (right) across ages. (**B**) Average expression of differentially expressed DNA methyltransferases across ages. (**C**) Median methylation levels of all DMRs within 100kb of signature genes for the specified cell types at each age. Boxes indicate gene sets corresponding to earlier ages of the same cell fate. (**D**) Normalized gene expression patterns, shown as z-scores, for the sets of signature genes used in panel C. (**E**) Size distribution of DMRs within each cluster. Bar: median; box: 25-75th quantile; whisker: 1.5 − inter-quartile range.

**Fig. S6.**
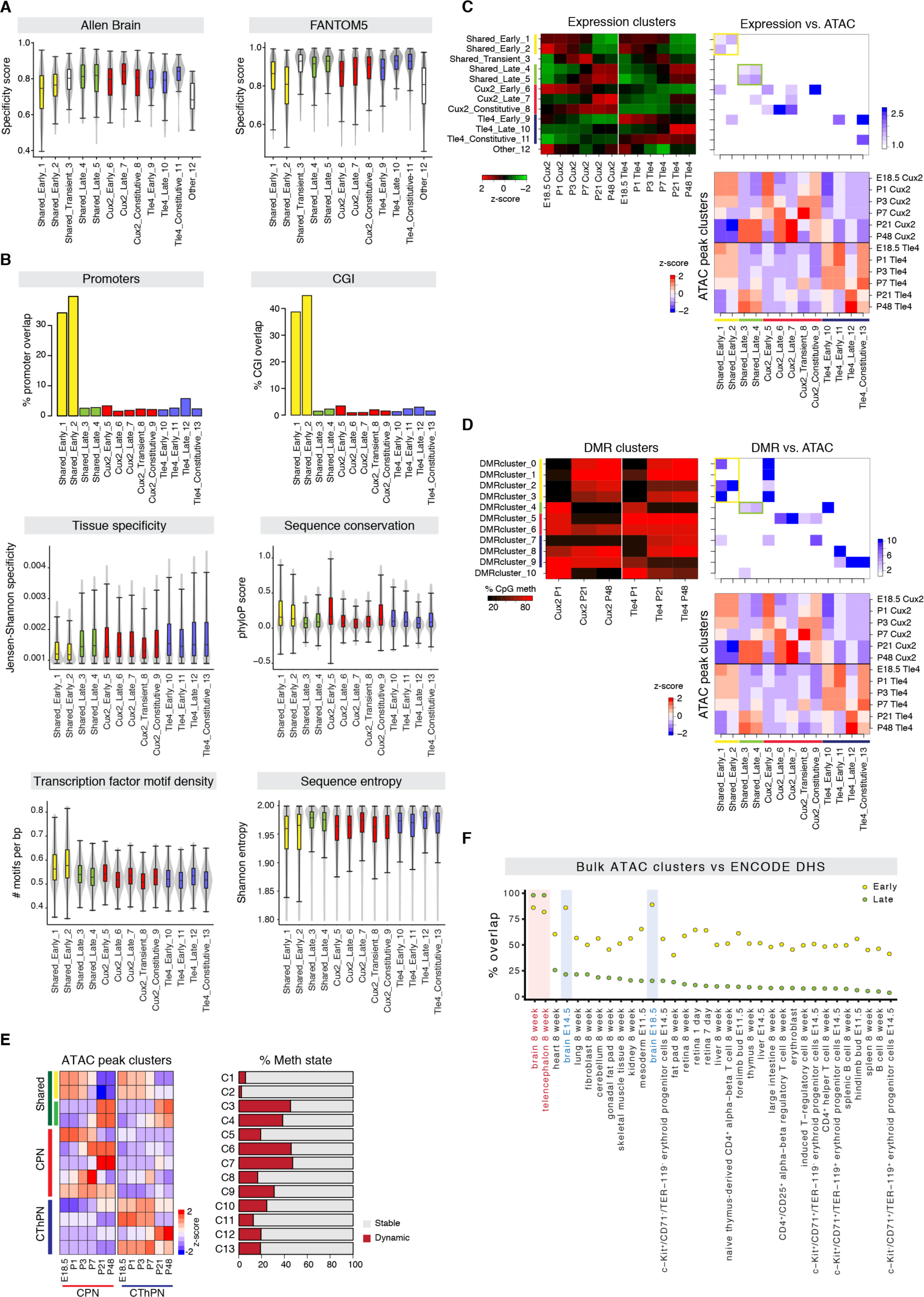
Characterization of RNAseq and ATACseq clusters by cluster. (**A**) Expression specificity of genes in each RNAseq cluster as in Fig. 2C, for each individual cluster. (**B**) Analyses of ATAC peak clusters as in Fig. 2E and H-K, for each individual cluster. (**C**) Correlation between individual RNAseq and ATAC clusters. (**D**) Correlation between individual DNAme and ATAC clusters. (**E**) Fraction of DMRs in the ATAC clusters that are dynamic vs. static over time as in as in Fig. 2F, for each ATAC cluster. (**F**) Overlap of early and late shared ATAC clusters with DNAse I hypersensitivity sites in a panel of tissues and cell types from the ENCODE database, by tissue.

**Fig. S7.**
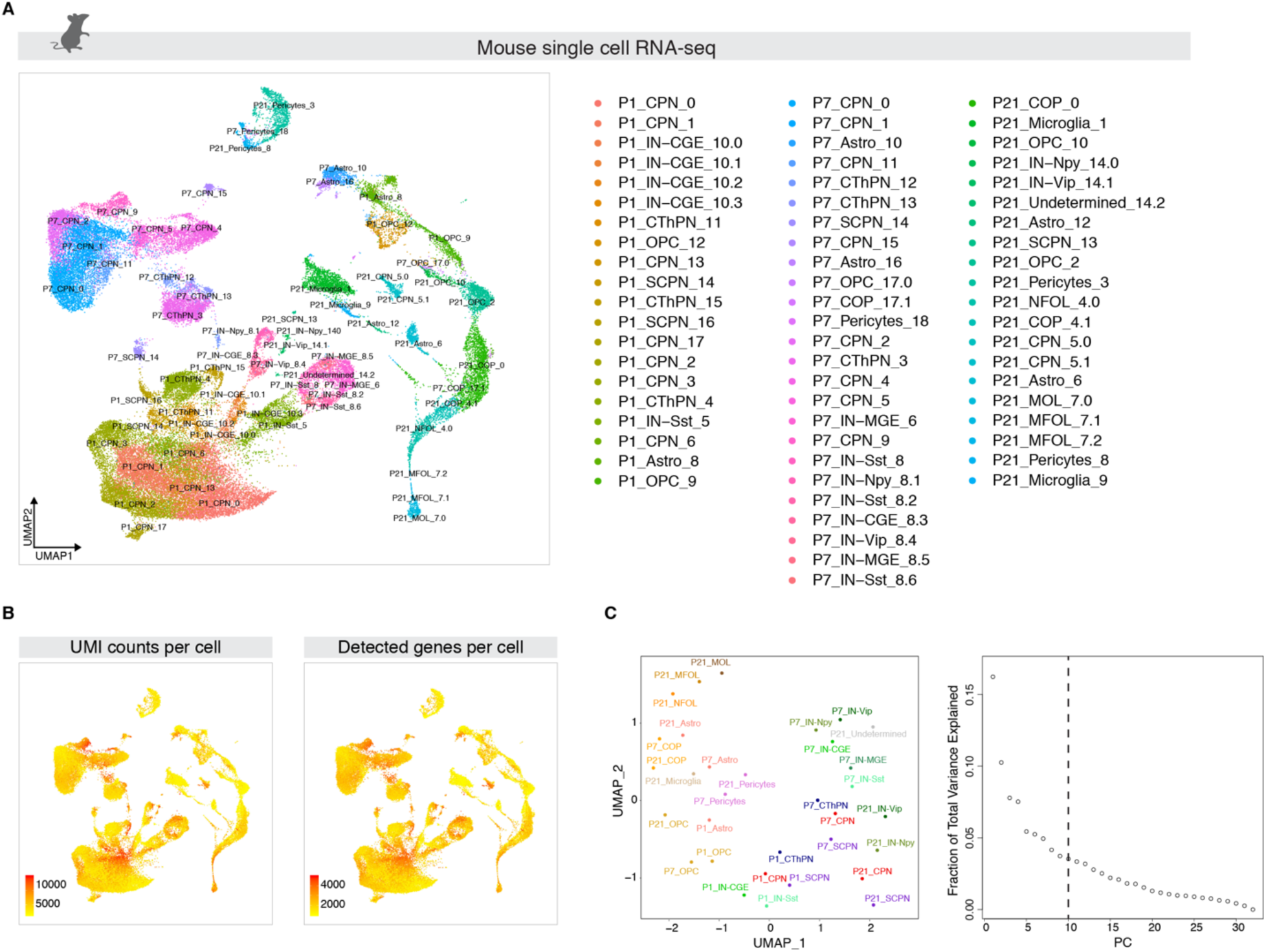
Characterization of mouse unfractionated single-cell RNAseq dataset. (**A**) UMAPs showing each cell cluster, broken down by age. (**B**) UMAPs showing number of UMIs and number of genes detected in each cell. (**C**) Similarity between cell clusters, as a UMAP representation of the top 10 principal components (PCs) after PCA computed on the average gene expression space.

**Fig. S8.**
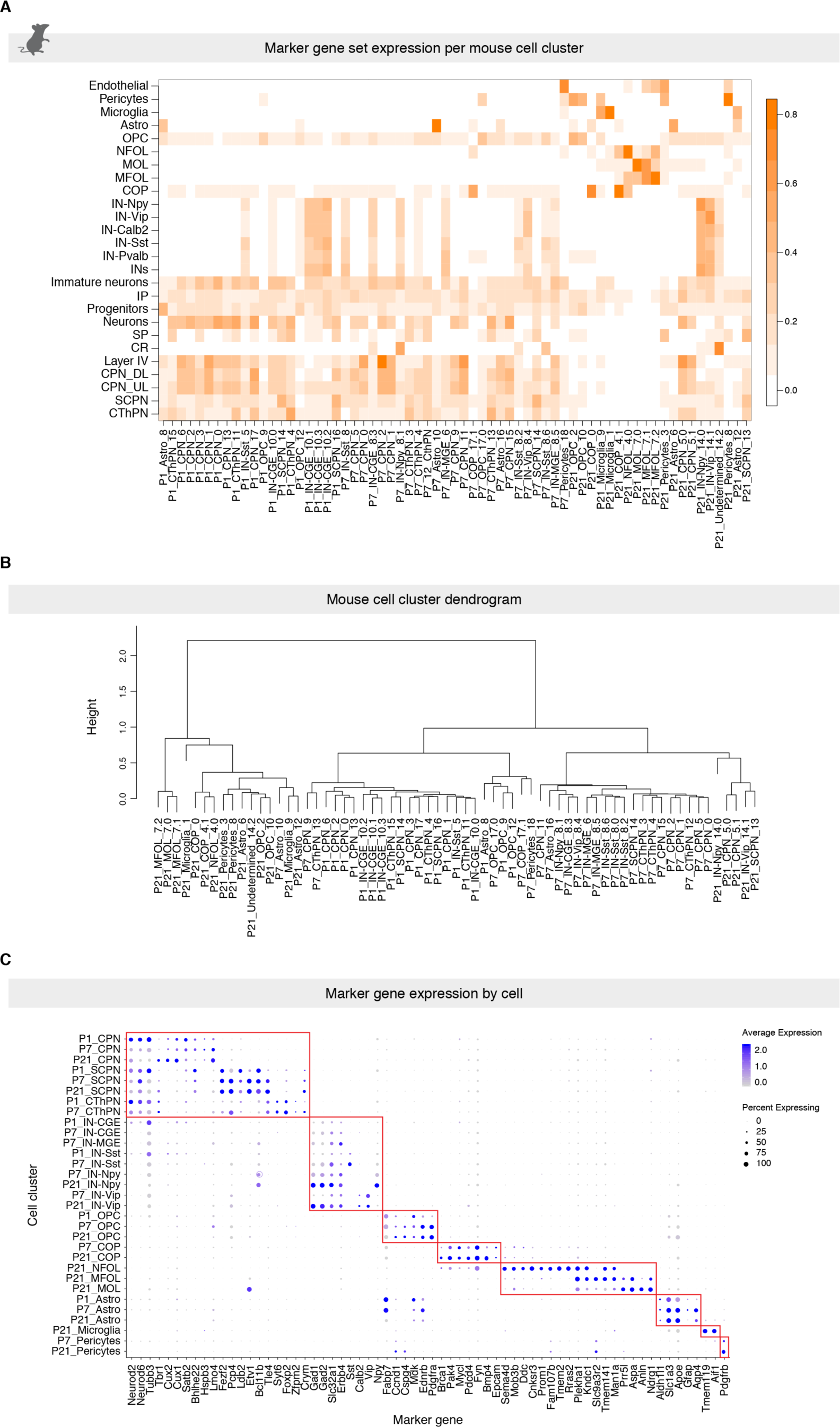
Characterization of mouse unfractionated single-cell RNAseq dataset. (**A**) Expression of gene sets representing combined expression of panels of known cell-type marker genes for each cell cluster. (**B**) Dendrogram representing similarity between cell clusters. (**C**) Average expression and percent of cells expressing for selected marker genes within cells of each assigned cell identity.

**Fig. S9.**
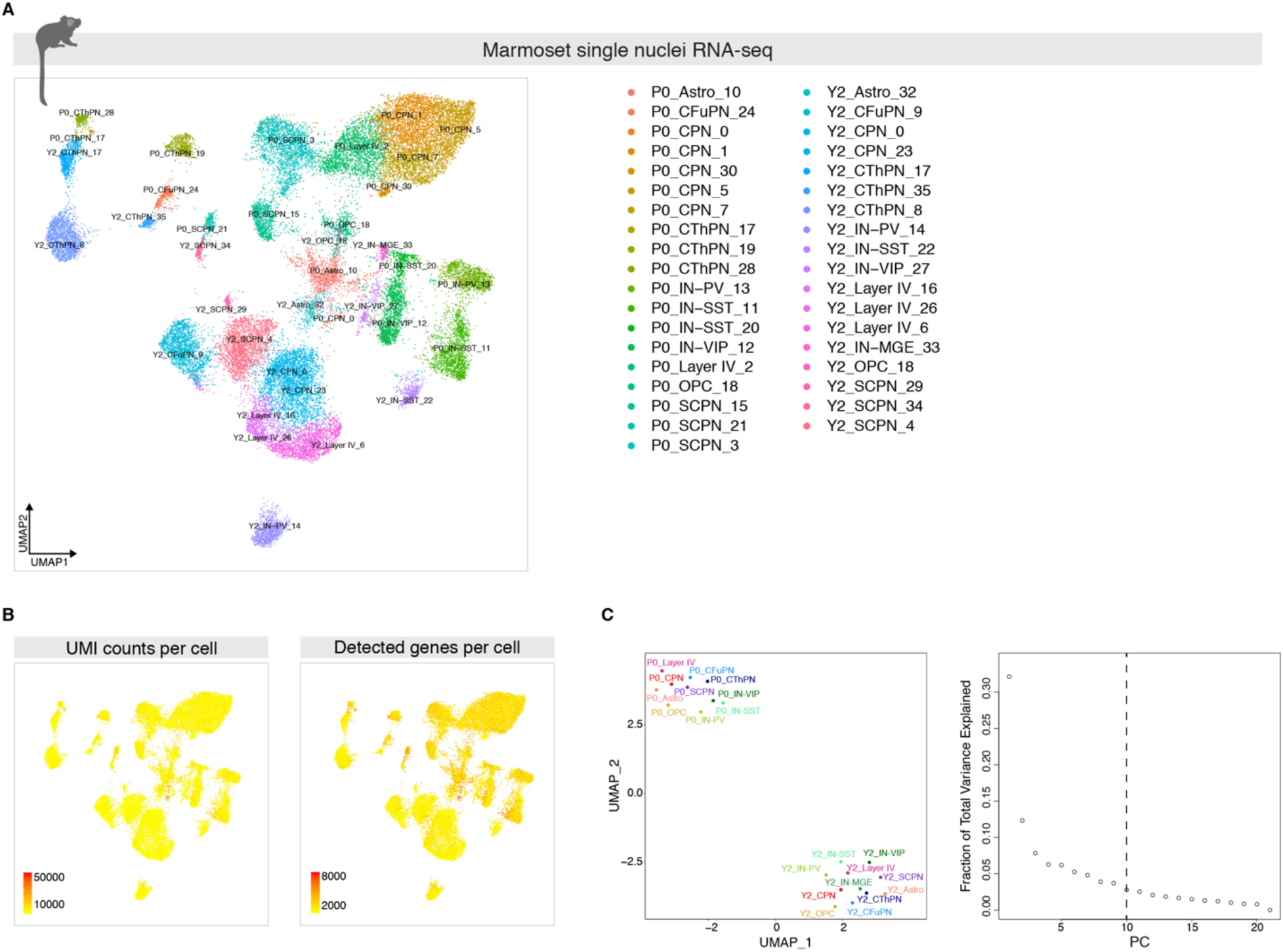
Characterization of marmoset single-cell RNAseq dataset. (**A**) UMAPs showing each cell cluster, broken down by age. (**B**) UMAPs showing number of UMIs and number of genes detected in each cell. (**C**) Similarity between cell clusters, as a UMAP representation of the top 10 principal components (PCs) after PCA computed on the average gene expression space.

**Fig. S10.**
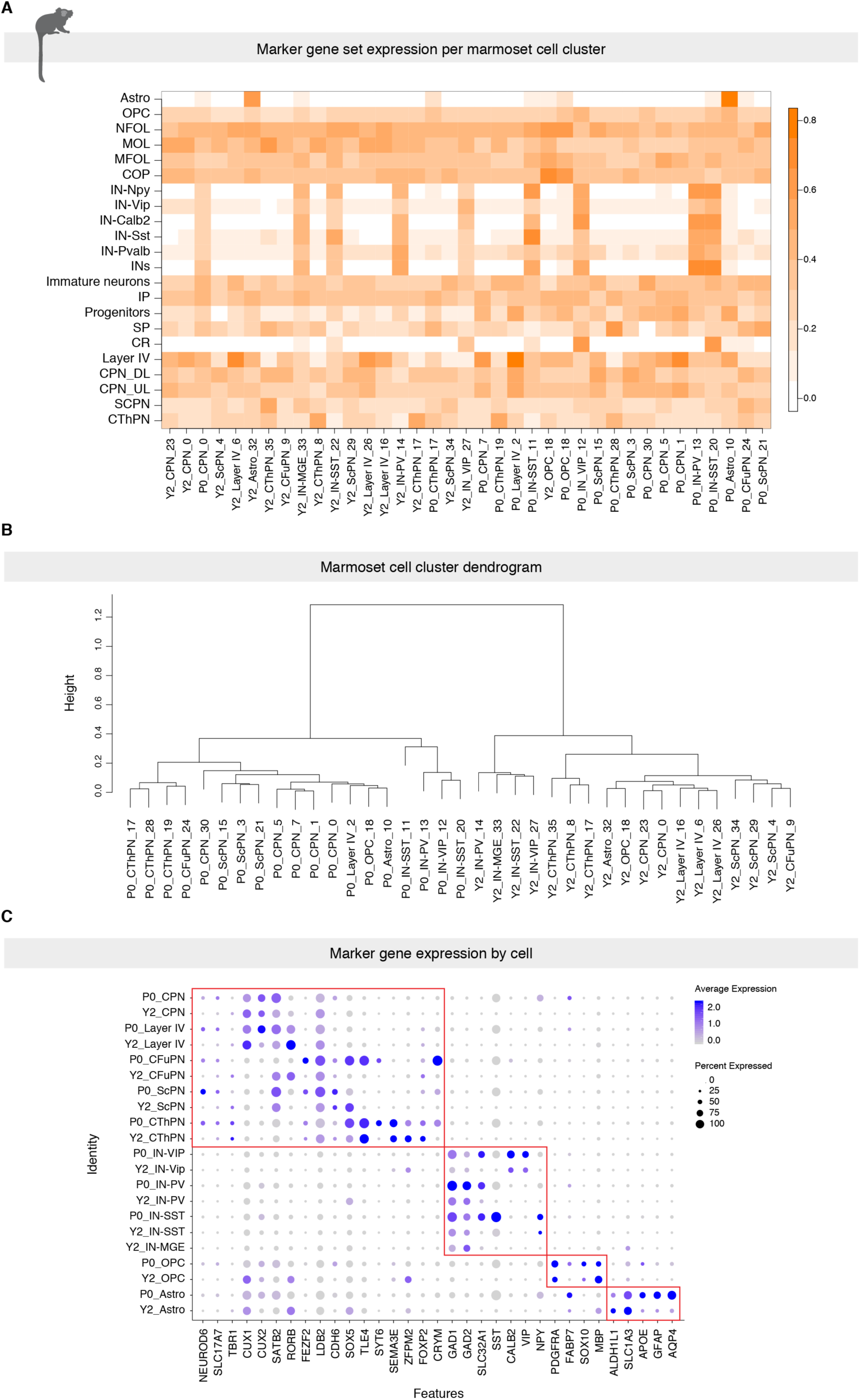
Characterization of marmoset single-cell RNAseq dataset. (**A**) Expression of gene sets representing combined expression of panels of known cell-type marker genes for each cell cluster. (**B**) Dendrogram representing similarity between cell clusters. (**C**) Average expression and percent of cells expressing for selected marker genes within cells of each assigned cell identity.

**Fig. S11.**
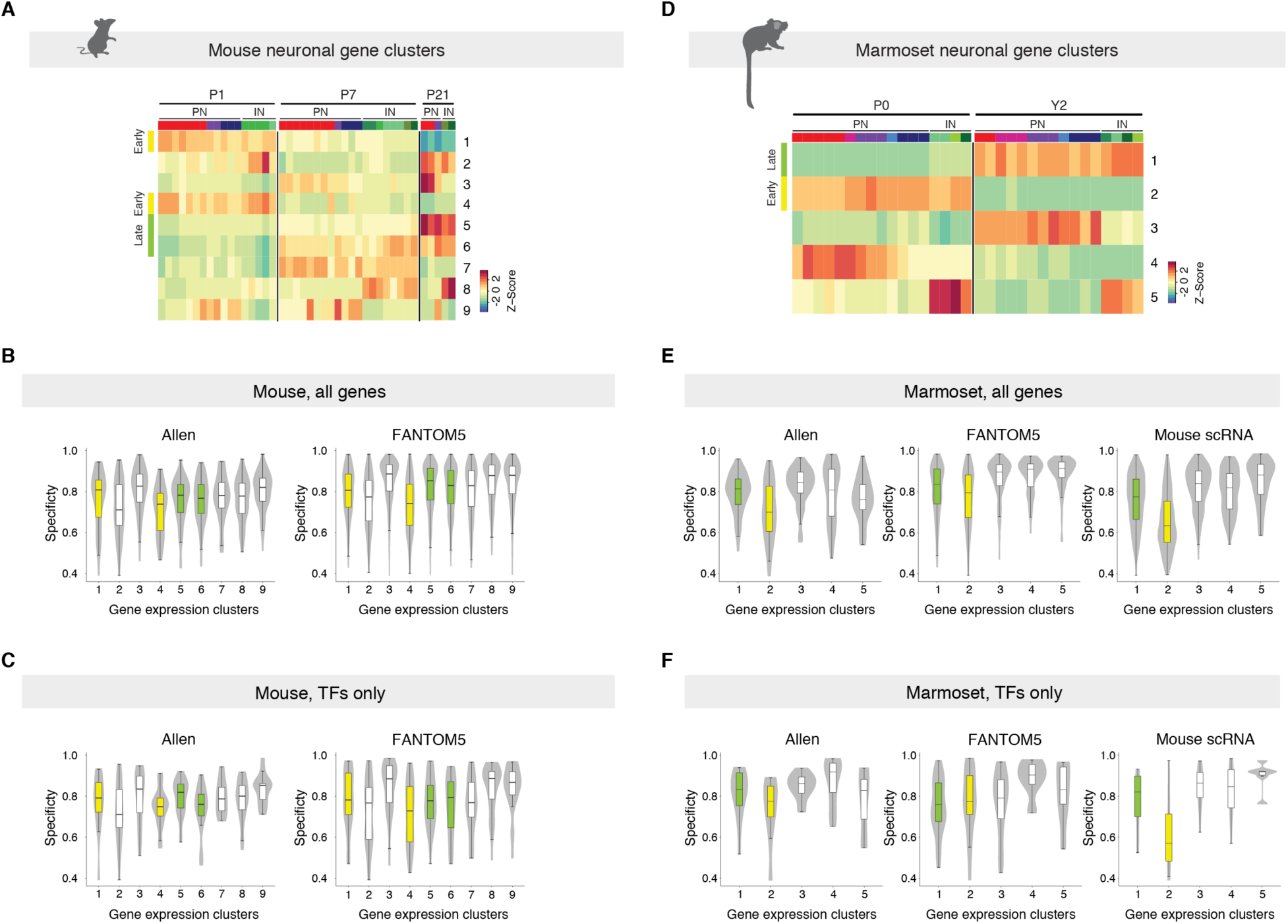
Gene cluster specificity. (**A**) Mouse neuronal gene clusters, repeated from Fig. 3 for ease of reference, in unsorted order. (**B**) Cell-type specificity of all genes in the Allen (left) and FANTOM5 (right) datasets, for each gene cluster. (**C**) Cell-type specificity of transcription factors (TFs) in the Allen (left) and FANTOM5 (right) datasets, for each gene cluster. (**D**) Marmoset neuronal gene clusters, repeated from Fig. 3 for ease of reference, in unsorted order. (**E**) Cell-type specificity of all genes in the Allen (left) and FANTOM5 (right) datasets, and in our mouse scRNA-seq dataset for each gene cluster. (**F**) Cell-type specificity of transcription factors in the Allen (left) and FANTOM5 (center) datasets, and in our mouse scRNA-seq dataset (right), for each gene cluster.

**Fig. S12.**
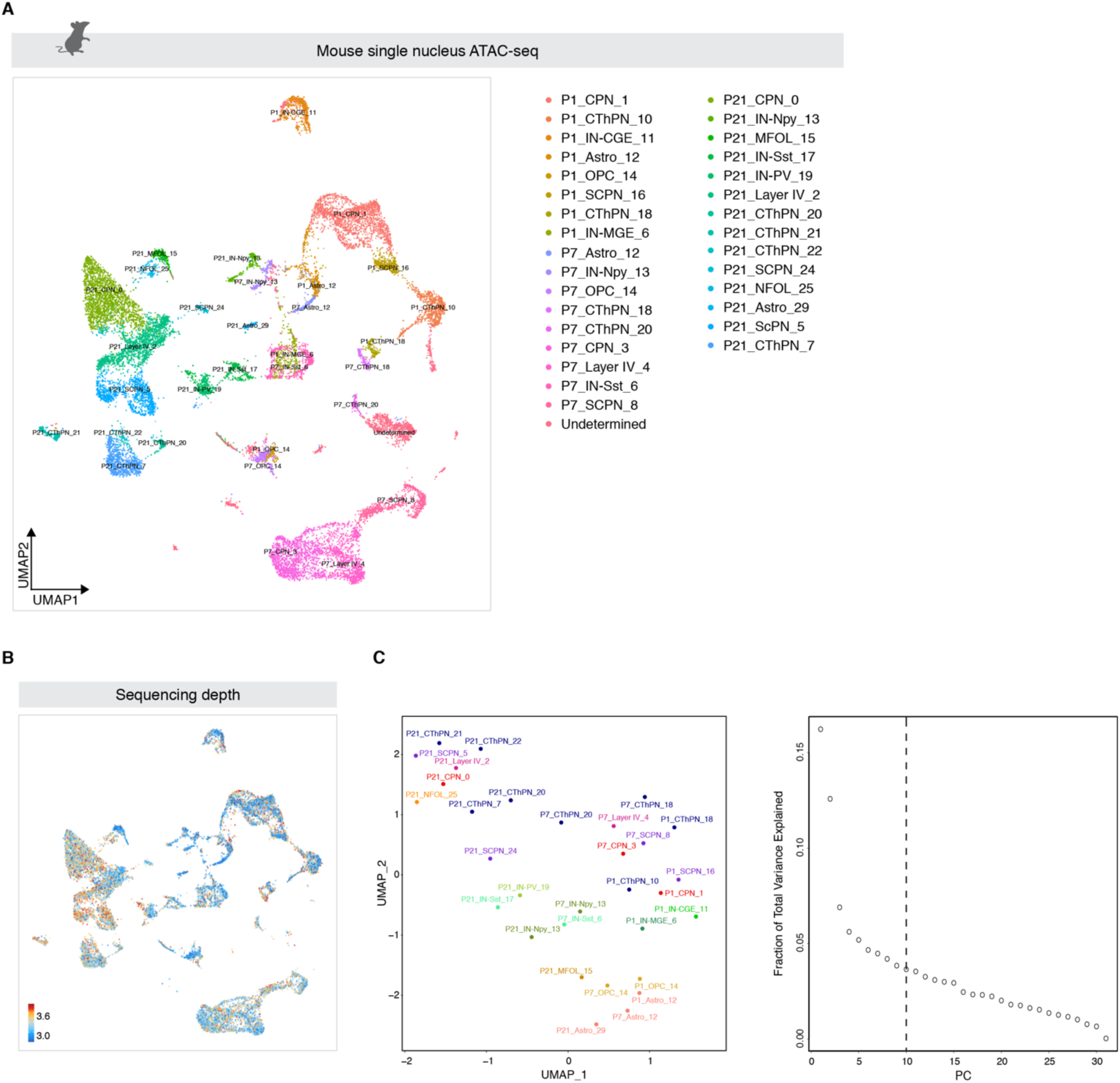
Characterization of mouse single-nucleus ATAC-seq dataset. (**A**) UMAPs showing each cell cluster, broken down by age. (**B**) UMAP showing depth of sequencing for each cell. (**C**) Similarity between cell clusters, as a UMAP representation of the top 10 principal components (PCs) after PCA computed on the average gene expression space.

**Fig. S13.**
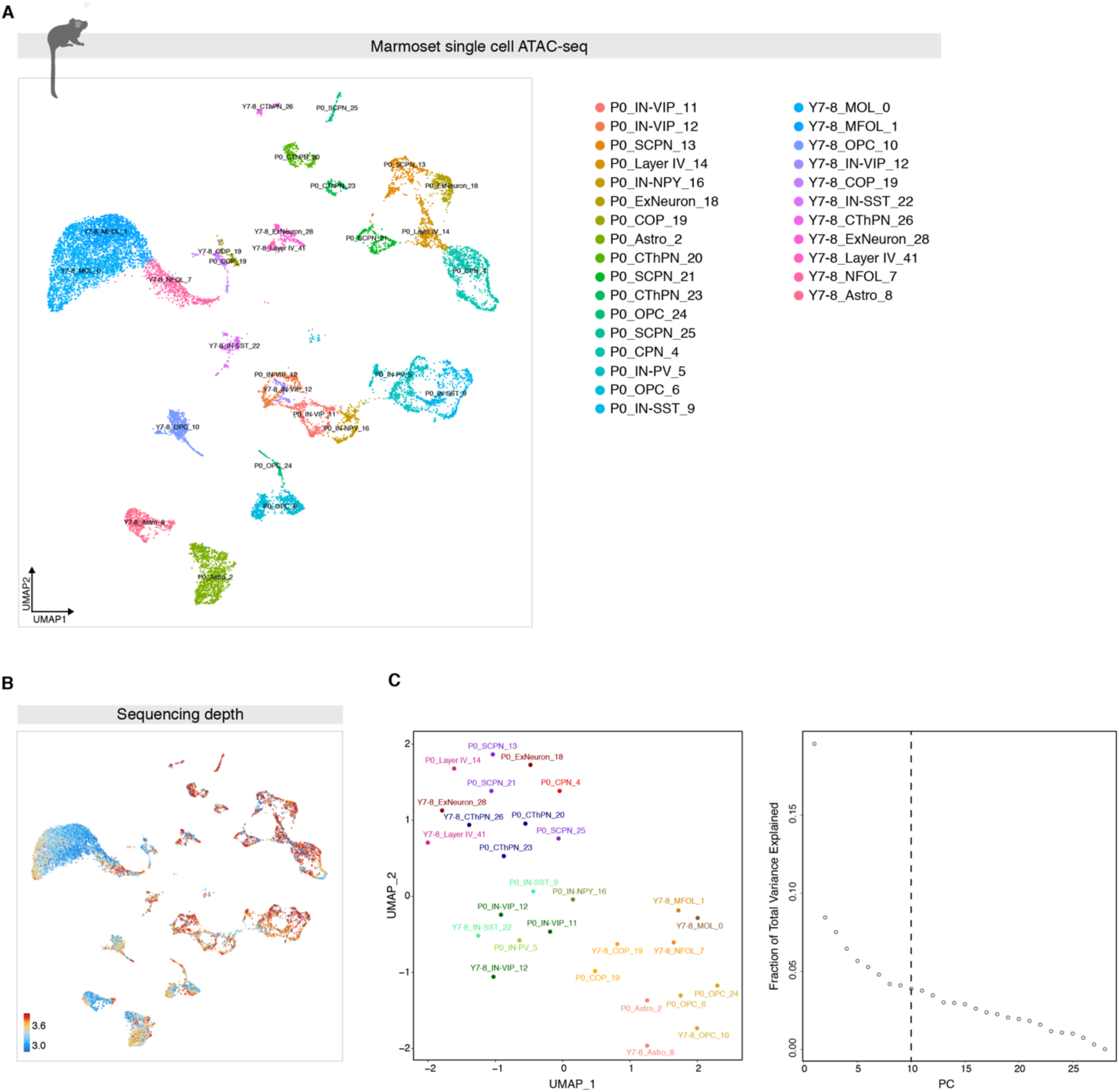
Characterization of marmoset single-cell ATAC-seq dataset. (**A**) UMAPs showing each cell cluster, broken down by age. (**B**) UMAP showing depth of sequencing for each cell. (**C**) Similarity between cell clusters, as a UMAP representation of the top 10 principal components (PCs) after PCA computed on the average gene expression space.

**Fig. S14.**
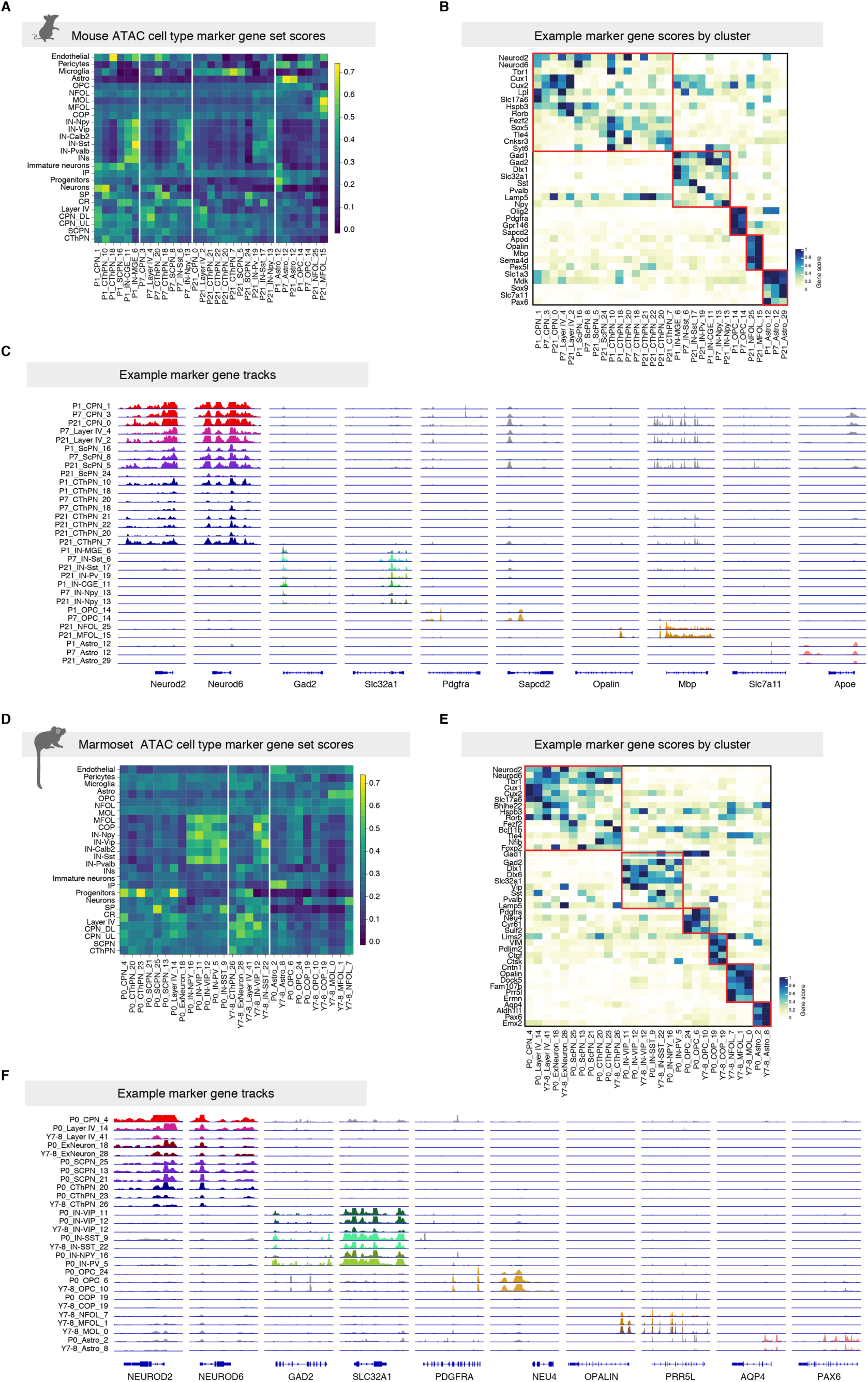
Cell-type assignments for scATAC datasets. (**A**) Average gene score for mouse assigned cell identities using the same panel of cell-type marker gene sets as in Fig. S8. (**B**) Gene score for example individual marker genes. (**C**) Mouse gene tracks for example marker genes across each assigned cell identity. (**D**) Average gene score for marmoset assigned cell identities using the same panels of cell-type marker gene sets as in Fig. S10. (**E**) Gene score for example individual marker genes. (**F**) Marmoset gene tracks for example marker genes across each assigned cell identity.

**Fig. S15.**
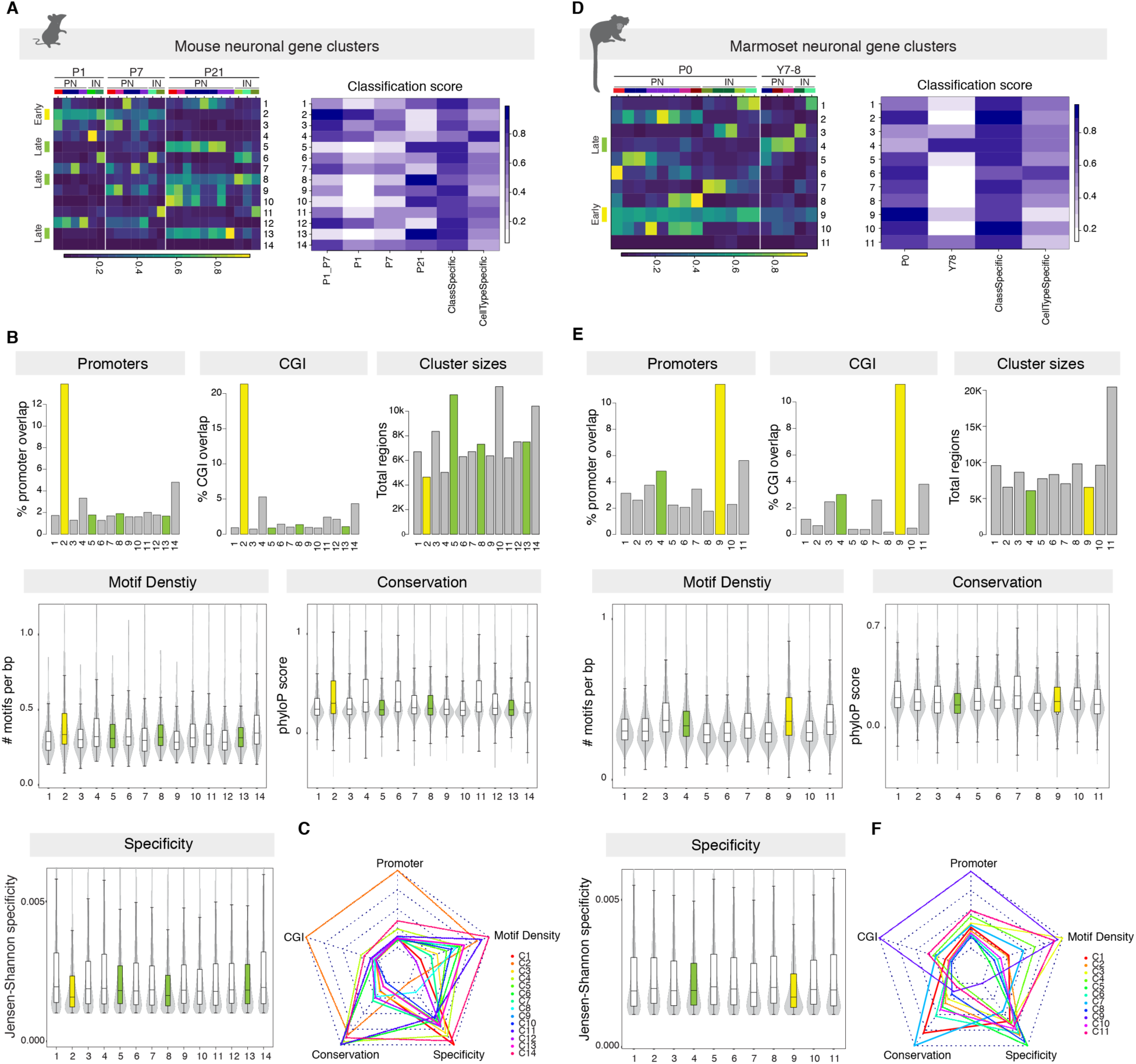
Specificity analyses of mouse and marmoset scATAC peak clusters. (**A**) Mouse ATAC peak clusters, reproduced from Fig. 4 for ease of reference, in unsorted order (left), and correlation with age vs. cell type for each cluster (right). (**B**) Mouse ATAC cluster characteristics as in Fig. 4, for each individual cluster, and number of regions per cluster (Cluster sizes). (**C**) Marmoset ATAC peak clusters, reproduced from Fig. 4 for ease of reference, in unsorted order (left), and correlation with age vs. cell type for each cluster (right). (**D**) Marmoset ATAC cluster characteristics as in Fig. 4, for each individual cluster, and number of regions per cluster (Cluster sizes).

**Fig. S16.**
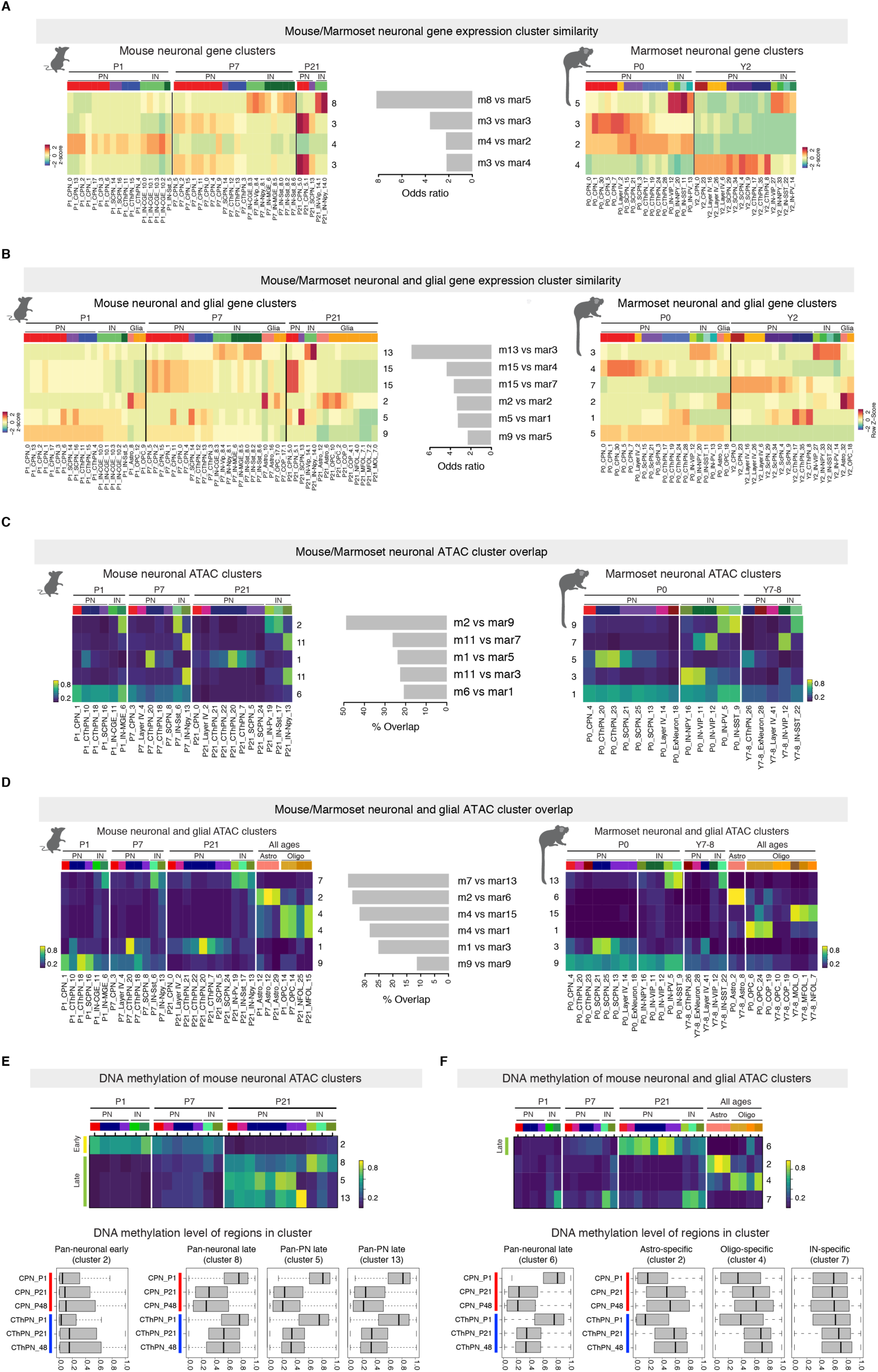
Correlation of mouse vs. marmoset gene and ATAC clusters. (**A**) Overlap between mouse and marmoset gene clusters for neuronal cell types only (as presented in text), showing the top 4 pairs. (**B**) Overlap between mouse and marmoset gene clusters for neuronal and glial cell types taken together (for comparison as a control), showing the top 6 pairs. (**C**) Overlap between mouse and marmoset ATAC clusters for neuronal cell types only, showing the top 5 pairs. (**D**) Overlap between mouse and marmoset ATAC clusters for neuronal and glial cell types taken together, showing the top 4 pairs. (**E**) Methylation of ATAC peaks in each mouse shared-developmental neuronal ATAC cluster in the mouse bulk DNAme dataset in Fig. 1. (**F**) Methylation of ATAC peaks in each mouse shared-developmental neuronal and glial ATAC cluster, determined from the mouse bulk WGBS dataset presented in Fig. 1.

## References

Akalin, A., Kormaksson, M., Li, S., Garrett-Bakelman, F.E., Figueroa, M.E., Melnick, A., and Mason, C.E. (2012). methylKit: a comprehensive R package for the analysis of genome-wide DNA methylation profiles. Genome Biol 13, R87.

Arlotta, P., Molyneaux, B.J., Chen, J., Inoue, J., Kominami, R., and Macklis, J.D. (2005). Neuronal subtype-specific genes that control corticospinal motor neuron development in vivo. Neuron 45, 207–221.

Ashburner, M., Ball, C.A., Blake, J.A., Botstein, D., Butler, H., Cherry, J.M., Davis, A.P., Dolinski, K., Dwight, S.S., Eppig, J.T., et al. (2000). Gene ontology: tool for the unification of biology. The Gene Ontology Consortium. Nat Genet 25, 25–29.

Baker, S.M., Rogerson, C., Hayes, A., Sharrocks, A.D., and Rattray, M. (2019). Classifying cells with Scasat, a single-cell ATAC-seq analysis tool. Nucleic Acids Res 47, e10.

Benjamini, Y., and Hochberg, Y. (1995). Controlling the False Discovery Rate: A Practical and Powerful Approach to Multiple Testing. Journal of the Royal Statistical Society, Series B 57, 289–300.

Bock, C., Beerman, I., Lien, W.H., Smith, Z.D., Gu, H., Boyle, P., Gnirke, A., Fuchs, E., Rossi, D.J., and Meissner, A. (2012). DNA methylation dynamics during in vivo differentiation of blood and skin stem cells. Mol Cell 47, 633–647.

Bolger, A.M., Lohse, M., and Usadel, B. (2014). Trimmomatic: a flexible trimmer for Illumina sequence data. Bioinformatics 30, 2114–2120.

Broad Institute of MIT and Harvard (2020). Picard. http://broadinstitute.github.io/picard.

Buenrostro, J.D., Giresi, P.G., Zaba, L.C., Chang, H.Y., and Greenleaf, W.J. (2013). Transposition of native chromatin for fast and sensitive epigenomic profiling of open chromatin, DNA-binding proteins and nucleosome position. Nat Methods 10, 1213–1218.

Butler, A., Hoffman, P., Smibert, P., Papalexi, E., and Satija, R. (2018). Integrating single-cell transcriptomic data across different conditions, technologies, and species. Nat Biotechnol 36, 411–420.

Cahoy, J.D., Emery, B., Kaushal, A., Foo, L.C., Zamanian, J.L., Christopherson, K.S., Xing, Y., Lubischer, J.L., Krieg, P.A., Krupenko, S.A., et al. (2008). A transcriptome database for astrocytes, neurons, and oligodendrocytes: a new resource for understanding brain development and function. The Journal of neuroscience : the official journal of the Society for Neuroscience 28, 264–278.

Cock, P.J., Antao, T., Chang, J.T., Chapman, B.A., Cox, C.J., Dalke, A., Friedberg, I., Hamelryck, T., Kauff, F., Wilczynski, B., et al. (2009). Biopython: freely available Python tools for computational molecular biology and bioinformatics. Bioinformatics 25, 1422–1423.

Corces, M.R., Buenrostro, J.D., Wu, B., Greenside, P.G., Chan, S.M., Koenig, J.L., Snyder, M.P., Pritchard, J.K., Kundaje, A., Greenleaf, W.J., et al. (2016). Lineage-specific and single-cell chromatin accessibility charts human hematopoiesis and leukemia evolution. Nat Genet 48, 1193–1203.

Cusanovich, D.A., Daza, R., Adey, A., Pliner, H.A., Christiansen, L., Gunderson, K.L., Steemers, F.J., Trapnell, C., and Shendure, J. (2015). Multiplex single cell profiling of chromatin accessibility by combinatorial cellular indexing. Science 348, 910–914.

Cusanovich, D.A., Hill, A.J., Aghamirzaie, D., Daza, R.M., Pliner, H.A., Berletch, J.B., Filippova, G.N., Huang, X., Christiansen, L., DeWitt, W.S., et al. (2018). A Single-Cell Atlas of In Vivo Mammalian Chromatin Accessibility. Cell 174, 1309–1324.e1318.

Darmanis, S., Sloan, S.A., Zhang, Y., Enge, M., Caneda, C., Shuer, L.M., Hayden Gephart, M.G., Barres, B.A., and Quake, S.R. (2015). A survey of human brain transcriptome diversity at the single cell level. Proc Natl Acad Sci U S A 112, 7285–7290.

Deaton, A.M., and Bird, A. (2011). CpG islands and the regulation of transcription. Genes Dev 25, 1010–1022.

Ehrlich, M., and Lacey, M. (2013). DNA methylation and differentiation: silencing, upregulation and modulation of gene expression. Epigenomics 5, 553–568.

Fame, R.M., MacDonald, J.L., and Macklis, J.D. (2011). Development, specification, and diversity of callosal projection neurons. Trends in Neurosciences 34, 41–50.

FANTOM Consortium and the RIKEN PMI and CLST (DGT), Forrest, A.R., Kawaji, H., Rehli, M., Baillie, J.K., de Hoon, M.J., Haberle, V., Lassmann, T., Kulakovskiy, I.V., Lizio, M., et al. (2014). A promoter-level mammalian expression atlas. Nature 507, 462–470.

Fogel, G.B., Weekes, D.G., Varga, G., Dow, E.R., Craven, A.M., Harlow, H.B., Su, E.W., Onyia, J.E., and Su, C. (2005). A statistical analysis of the TRANSFAC database. Biosystems 81, 137–154.

Franco, S.J., Gil-Sanz, C., Martinez-Garay, I., Espinosa, A., Harkins-Perry, S.R., Ramos, C., and Muller, U. (2012). Fate-restricted neural progenitors in the mammalian cerebral cortex. Science 337, 746–749.

Frazer, S., Prados, J., Niquille, M., Cadilhac, C., Markopoulos, F., Gomez, L., Tomasello, U., Telley, L., Holtmaat, A., Jabaudon, D., et al. (2017). Transcriptomic and anatomic parcellation of 5-HT3AR expressing cortical interneuron subtypes revealed by single-cell RNA sequencing. Nat Commun 8, 14219.

Galazo, M.J., Emsley, J.G., and Macklis, J.D. (2016). Corticothalamic Projection Neuron Development beyond Subtype Specification: Fog2 and Intersectional Controls Regulate Intraclass Neuronal Diversity. Neuron 91, 90–106.

Gil-Sanz, C., Espinosa, A., Fregoso, S.P., Bluske, K.K., Cunningham, C.L., Martinez-Garay, I., Zeng, H., Franco, S.J., and Müller, U. (2015). Lineage Tracing Using Cux2-Cre and Cux2-CreERT2 Mice. Neuron 86, 1091–1099.

Grant, C.E., Bailey, T.L., and Noble, W.S. (2011). FIMO: scanning for occurrences of a given motif. Bioinformatics 27, 1017–1018.

Gray, L.T., Yao, Z., Nguyen, T.N., Kim, T.K., Zeng, H., and Tasic, B. (2017). Layer-specific chromatin accessibility landscapes reveal regulatory networks in adult mouse visual cortex. Elife 6.

Guibert, S., and Weber, M. (2013). Functions of DNA methylation and hydroxymethylation in mammalian development. Curr Top Dev Biol 104, 47–83.

Hill, A. (2019). Dimensionality Reduction for scATAC Data. http://andrewjohnhill.com/blog/2019/05/06/dimensionality-reduction-for-scatac-data/.

Hodge, R.D., Bakken, T.E., Miller, J.A., Smith, K.A., Barkan, E.R., Graybuck, L.T., Close, J.L., Long, B., Johansen, N., Penn, O., et al. (2019). Conserved cell types with divergent features in human versus mouse cortex. Nature 573, 61–68.

Hon, G.C., Rajagopal, N., Shen, Y., McCleary, D.F., Yue, F., Dang, M.D., and Ren, B. (2013). Epigenetic memory at embryonic enhancers identified in DNA methylation maps from adult mouse tissues. Nat Genet 45, 1198–1206.

Jolma, A., Yan, J., Whitington, T., Toivonen, J., Nitta, K.R., Rastas, P., Morgunova, E., Enge, M., Taipale, M., Wei, G., et al. (2013). DNA-binding specificities of human transcription factors. Cell 152, 327–339.

Kent, W.J., Sugnet, C.W., Furey, T.S., Roskin, K.M., Pringle, T.H., Zahler, A.M., and Haussler, D. (2002). The human genome browser at UCSC. Genome Res 12, 996–1006.

Kim, D., Pertea, G., Trapnell, C., Pimentel, H., Kelley, R., and Salzberg, S.L. (2013). TopHat2: accurate alignment of transcriptomes in the presence of insertions, deletions and gene fusions. Genome Biol 14, R36.

Kryuchkova-Mostacci, N., and Robinson-Rechavi, M. (2017). A benchmark of gene expression tissue-specificity metrics. Brief Bioinform 18, 205–214.

Lafave et al., in press (2020). Cancer Cell.

Lake, B.B., Ai, R., Kaeser, G.E., Salathia, N.S., Yung, Y.C., Liu, R., Wildberg, A., Gao, D., Fung, H.L., Chen, S., et al. (2016). Neuronal subtypes and diversity revealed by single-nucleus RNA sequencing of the human brain. Science 352, 1586–1590.

Lake, B.B., Chen, S., Sos, B.C., Fan, J., Kaeser, G.E., Yung, Y.C., Duong, T.E., Gao, D., Chun, J., Kharchenko, P.V., et al. (2018). Integrative single-cell analysis of transcriptional and epigenetic states in the human adult brain. Nat Biotechnol 36, 70–80.

Langmead, B., and Salzberg, S.L. (2012). Fast gapped-read alignment with Bowtie 2. Nat Methods 9, 357–359.

Lawrence, M., Huber, W., Pages, H., Aboyoun, P., Carlson, M., Gentleman, R., Morgan, M.T., and Carey, V.J. (2013). Software for computing and annotating genomic ranges. PLoS Comput Biol 9, e1003118.

Lein, E.S., Hawrylycz, M.J., Ao, N., Ayres, M., Bensinger, A., Bernard, A., Boe, A.F., Boguski, M.S., Brockway, K.S., Byrnes, E.J., et al. (2007). Genome-wide atlas of gene expression in the adult mouse brain. Nature 445, 168–176.

Li, H., Handsaker, B., Wysoker, A., Fennell, T., Ruan, J., Homer, N., Marth, G., Abecasis, G., and Durbin, R. (2009). The Sequence Alignment/Map format and SAMtools. Bioinformatics 25, 2078–2079.

Li, Q., Brown, J.B., Huang, H., and Bickel, P.J. (2011). Measuring reproducibility of high-throughput experiments. Ann Appl Stat 5, 1752–1779.

Lodato, S., and Arlotta, P. (2015). Generating neuronal diversity in the mammalian cerebral cortex. Annu Rev Cell Dev Biol 31, 699–720.

Love, M.I., Huber, W., and Anders, S. (2014). Moderated estimation of fold change and dispersion for RNA-seq data with DESeq2. Genome Biol 15, 550.

Luo, C., Keown, C.L., Kurihara, L., Zhou, J., He, Y., Li, J., Castanon, R., Lucero, J., Nery, J.R., Sandoval, J.P., et al. (2017). Single-cell methylomes identify neuronal subtypes and regulatory elements in mammalian cortex. Science 357, 600–604.

Ma, S., de la Fuente Revenga, M., Sun, Z., Sun, C., Murphy, T.W., Xie, H., González-Maeso, J., and Lu, C. (2018). Cell-type-specific brain methylomes profiled via ultralow-input microfluidics. Nat Biomed Eng 2, 183–194.

Madisen, L., Zwingman, T.A., Sunkin, S.M., Oh, S.W., Zariwala, H.A., Gu, H., Ng, L.L., Palmiter, R.D., Hawrylycz, M.J., Jones, A.R., et al. (2010). A robust and high-throughput Cre reporting and characterization system for the whole mouse brain. Nat Neurosci 13, 133–140.

Maechler, M.R. Peter; Struyf, Anja; Hubert, Mia; Hornik, Kurt (2019). cluster: Cluster Analysis Basics and Extensions.

Matho, K.S., Huilgol, D., Galbavy, W., Kim, G., He, M., An, X., Lu, J., Wu, P., Di Bella, D.J., Shetty, A.S., et al. (2020). Genetic dissection of glutamatergic neuron subpopulations and developmental trajectories in the cerebral cortex. bioRxiv.

McKinley, K.L., Castillo-Azofeifa, D., and Klein, O.D. (2020). Tools and Concepts for Interrogating and Defining Cellular Identity. Cell Stem Cell 26, 632–656.

Mi, H., Muruganujan, A., Ebert, D., Huang, X., and Thomas, P.D. (2019). PANTHER version 14: more genomes, a new PANTHER GO-slim and improvements in enrichment analysis tools. Nucleic Acids Res 47, D419–d426.

Mo, A., Mukamel, E.A., Davis, F.P., Luo, C., Henry, G.L., Picard, S., Urich, M.A., Nery, J.R., Sejnowski, T.J., Lister, R., et al. (2015). Epigenomic Signatures of Neuronal Diversity in the Mammalian Brain. Neuron 86, 1369–1384.

Molyneaux, B.J., Goff, L.A., Brettler, A.C., Chen, H.H., Brown, J.R., Hrvatin, S., Rinn, J.L., and Arlotta, P. (2015). DeCoN: genome-wide analysis of in vivo transcriptional dynamics during pyramidal neuron fate selection in neocortex. Neuron 85, 275–288.

Natarajan, A., Yardimci, G.G., Sheffield, N.C., Crawford, G.E., and Ohler, U. (2012). Predicting cell-type-specific gene expression from regions of open chromatin. Genome Res 22, 1711–1722.

NIH Neuroscience Blueprint Cre Driver Network (2009). Cre recombinase-expressing mice generated for the NIH Neuroscience Blueprint Cre Driver Network (MGI Direct Data Submission).

Nott, A., Holtman, I.R., Coufal, N.G., Schlachetzki, J.C.M., Yu, M., Hu, R., Han, C.Z., Pena, M., Xiao, J., Wu, Y., et al. (2019). Brain cell type-specific enhancer-promoter interactome maps and disease-risk association. Science 366, 1134–1139.

Park, Y., and Wu, H. (2016). Differential methylation analysis for BS-seq data under general experimental design. Bioinformatics 32, 1446–1453.

Pliner, H.A., Packer, J.S., McFaline-Figueroa, J.L., Cusanovich, D.A., Daza, R.M., Aghamirzaie, D., Srivatsan, S., Qiu, X., Jackson, D., Minkina, A., et al. (2018). Cicero Predicts cis-Regulatory DNA Interactions from Single-Cell Chromatin Accessibility Data. Mol Cell 71, 858–871.e858.

Pollard, K.S., Hubisz, M.J., Rosenbloom, K.R., and Siepel, A. (2010). Detection of nonneutral substitution rates on mammalian phylogenies. Genome Res 20, 110–121.

Ravasi, T., Suzuki, H., Cannistraci, C.V., Katayama, S., Bajic, V.B., Tan, K., Akalin, A., Schmeier, S., Kanamori-Katayama, M., Bertin, N., et al. (2010). An atlas of combinatorial transcriptional regulation in mouse and man. Cell 140, 744–752.

Ross-Innes, C.S., Stark, R., Teschendorff, A.E., Holmes, K.A., Ali, H.R., Dunning, M.J., Brown, G.D., Gojis, O., Ellis, I.O., Green, A.R., et al. (2012). Differential oestrogen receptor binding is associated with clinical outcome in breast cancer. Nature 481, 389–393.

Song, L., Zhang, Z., Grasfeder, L.L., Boyle, A.P., Giresi, P.G., Lee, B.K., Sheffield, N.C., Graf, S., Huss, M., Keefe, D., et al. (2011). Open chromatin defined by DNaseI and FAIRE identifies regulatory elements that shape cell-type identity. Genome Res 21, 1757–1767.

Stark, R., and Brown, G. (2011). DiffBind: differential binding analysis of ChIP-Seq peak data (R package).

Stuart, T., Butler, A., Hoffman, P., Hafemeister, C., Papalexi, E., Mauck, W.M., 3rd, Hao, Y., Stoeckius, M., Smibert, P., and Satija, R. (2019). Comprehensive Integration of Single-Cell Data. Cell 177, 1888–1902.e1821.

Sun, D., Xi, Y., Rodriguez, B., Park, H.J., Tong, P., Meong, M., Goodell, M.A., and Li, W. (2014). MOABS: model based analysis of bisulfite sequencing data. Genome Biol 15, R38.

Tasic, B., Menon, V., Nguyen, T.N., Kim, T.K., Jarsky, T., Yao, Z., Levi, B., Gray, L.T., Sorensen, S.A., Dolbeare, T., et al. (2016). Adult mouse cortical cell taxonomy revealed by single cell transcriptomics. Nat Neurosci 19, 335–346.

The Gene Ontology Consortium (2019). The Gene Ontology Resource: 20 years and still GOing strong. Nucleic Acids Res 47, D330–d338.

Thompson, C.L., Ng, L., Menon, V., Martinez, S., Lee, C.K., Glattfelder, K., Sunkin, S.M., Henry, A., Lau, C., Dang, C., et al. (2014). A high-resolution spatiotemporal atlas of gene expression of the developing mouse brain. Neuron 83, 309–323.

Tibshirani, R., Walther, G., and Hastie, T. (2001). Estimating the number of clusters in a data set via the gap statistic. Journal of the Royal Statistical Society: Series B (Statistical Methodology) 63, 411–423.

Trapnell, C., Hendrickson, D.G., Sauvageau, M., Goff, L., Rinn, J.L., and Pachter, L. (2013). Differential analysis of gene regulation at transcript resolution with RNA-seq. Nat Biotechnol 31, 46–53.

Xi, Y., and Li, W. (2009). BSMAP: whole genome bisulfite sequence MAPping program. BMC Bioinformatics 10, 232.

Yao, Z., Liu, H., Xie, F., Fischer, S., Booeshaghi, A.S., Adkins, R.S., Aldridge, A.I., Ament, S.A., Pinto-Duarte, A., Bartlett, A., et al. (2020). An integrated transcriptomic and epigenomic atlas of mouse primary motor cortex cell types. bioRxiv.

Yue, F., Cheng, Y., Breschi, A., Vierstra, J., Wu, W., Ryba, T., Sandstrom, R., Ma, Z., Davis, C., Pope, B.D., et al. (2014). A comparative encyclopedia of DNA elements in the mouse genome. Nature 515, 355–364.

Zeisel, A., Munoz-Manchado, A.B., Codeluppi, S., Lonnerberg, P., La Manno, G., Jureus, A., Marques, S., Munguba, H., He, L., Betsholtz, C., et al. (2015). Brain structure. Cell types in the mouse cortex and hippocampus revealed by single-cell RNA-seq. Science 347, 1138–1142.

Zhang, Y., Chen, K., Sloan, S.A., Bennett, M.L., Scholze, A.R., O’Keeffe, S., Phatnani, H.P., Guarnieri, P., Caneda, C., Ruderisch, N., et al. (2014). An RNA-sequencing transcriptome and splicing database of glia, neurons, and vascular cells of the cerebral cortex. J Neurosci 34, 11929–11947.

Zhu, L.J. (2013). Integrative Analysis of ChIP-Chip and ChIP-Seq Dataset. In Tiling Arrays: Methods and Protocols, T.-L. Lee, and A.C. Shui Luk, eds. (Totowa, NJ: Humana Press), pp. 105–124.

Zhu, L.J., Gazin, C., Lawson, N.D., Pages, H., Lin, S.M., Lapointe, D.S., and Green, M.R. (2010). ChIPpeakAnno: a Bioconductor package to annotate ChIP-seq and ChIP-chip data. BMC Bioinformatics 11, 237.

Ziller, M.J., Gu, H., Muller, F., Donaghey, J., Tsai, L.T., Kohlbacher, O., De Jager, P.L., Rosen, E.D., Bennett, D.A., Bernstein, B.E., et al. (2013). Charting a dynamic DNA methylation landscape of the human genome. Nature 500, 477–481.

